# Mountain stoneflies may tolerate warming streams: evidence from organismal physiology and gene expression

**DOI:** 10.1101/2019.12.16.878926

**Authors:** Scott Hotaling, Alisha A. Shah, Kerry L. McGowan, Lusha M. Tronstad, J. Joseph Giersch, Debra S. Finn, H. Arthur Woods, Michael E. Dillon, Joanna L. Kelley

## Abstract

Rapid glacier recession is altering the physical conditions of headwater streams. Stream temperatures are predicted to rise and become increasingly variable, putting entire meltwater-associated biological communities at risk of extinction. Thus, there is a pressing need to understand how thermal stress affects mountain stream insects, particularly where glaciers are likely to vanish on contemporary timescales. In this study, we tested the critical thermal maximum (CT_MAX_) of stonefly nymphs representing multiple species and a range of thermal regimes in the high Rocky Mountains, USA. We then collected RNA-sequencing data to assess how organismal thermal stress translated to the cellular level. Our focal species included the meltwater stonefly, *Lednia tumana*, which was recently listed under the U.S. Endangered Species Act due to climate-induced habitat loss. For all study species, critical thermal maxima (CT_MAX_ > 20°C) far exceeded the stream temperatures mountain stoneflies experience (< 10°C). Moreover, while evidence for a cellular stress response was present, we also observed constitutive expression of genes encoding proteins known to underlie thermal stress (i.e., heat shock proteins) even at low temperatures that reflected natural conditions. We show that high-elevation aquatic insects may not be physiologically threatened by short-term exposure to warm temperatures and that longer term physiological responses or biotic factors (e.g., competition) may better explain their extreme distributions.

## Introduction

Predicting how species will respond to climate change is a central goal of contemporary ecology (Araújo & New, 2007, Urban *et al*., 2016). This goal is difficult to achieve, however, because at a minimum it requires knowledge of extant distributions, physiological limits, and future conditions in relevant habitats. Mountain streams around the world are being transformed by climate change, primarily through rapid recession of glaciers and perennial snowfields (Hotaling *et al*., 2017). Warmer air temperatures are predicted to cause loss of permanent snow and ice, drive generally higher, more variable stream temperatures, and eventually lower flows in meltwater-dominated catchments (Huss & Hock, 2018, Jones *et al*., 2014). Rapid contemporary warming has already been observed in the European Alps, with streams warming at a mean rate of 2.5°C per decade (Niedrist & Füreder, 2020). Expected ecological responses include a reduction of biodiversity in headwater streams across multiple levels of biological organization and taxonomic groups (Bálint *et al*., 2011, Finn *et al*., 2013, Giersch *et al*., 2017, Hotaling *et al*., 2019a, Jordan *et al*., 2016). Considerable attention has been devoted to potential losses of aquatic insect diversity (e.g., Jacobsen *et al*., 2012). However, the specific mechanisms underlying physiological limits in alpine stream insects remain unknown. This knowledge gap is particularly important in light of the widely held assumption that aquatic insects living at high-elevations are cold-adapted stenotherms that will not tolerate warming streams (Giersch *et al*., 2015, Jacobsen *et al*., 2012). Recent evidence that the thermal maxima of high-elevation stream taxa can exceed maximum water temperatures (e.g., Shah *et al*., 2017b), that spring-dwelling cold stenotherms exhibit little variability in heat shock protein (HSP) expression across temperatures (e.g., Ebner *et al*., 2019), and that meltwater-associated invertebrate communities persist despite widespread deglaciation (Muhlfeld *et al*., 2020) all challenge this assumption, raising new questions about whether climate warming directly threatens headwater biodiversity. To better understand the degree to which headwater species can tolerate warming, links between relevant traits at the organismal (thermal stress) and cellular (e.g., gene expression) level are needed.

As ectotherms, insect body temperatures depend strongly on their external environment. Insects are therefore threatened by rising global temperatures, and recent studies have documented declines in their diversity (Lister & Garcia, 2018, Sánchez-Bayo & Wyckhuys, 2019). The effects of temperature on ectotherm performance and survival, however, are complex. Ectotherms may respond to stressful temperatures through plasticity or acclimatization (Seebacher *et al*., 2015), the evolution of higher thermal limits (Angilletta Jr *et al*., 2007), or behavioral thermoregulation (Kearney *et al*., 2009). Temperature can also affect organismal distributions indirectly. For instance, changing temperatures can alter ratios of oxygen supply and demand (Pörtner *et al*., 2007, Verberk *et al*., 2016b). Extreme temperatures can also provide natural buffering against invasions by competitors or predators (Isaak *et al*., 2015). Thus, temperature likely shapes both the evolution of aquatic insect physiology as well as local networks of biotic interactions (Shah *et al*., 2020). To understand the relationship between temperature and ectotherm tolerance, trait-based approaches (e.g., measuring upper thermal tolerance) can be informative. However, a focus on physiological traits at the whole-organism level may overlook other key aspects of a species’ potential for response, perhaps limiting predictions of whether species can evolve in response to changing thermal regimes (Chown *et al*., 2010) or tolerate them *in situ* via plasticity. Thus, there is a need to connect traits from cellular to organismal levels and consider findings holistically.

Due to the high heat capacity of water, stream temperatures are less thermally variable than air. However, a surprising amount of variation still exists in streams due to many factors, including latitude, elevation, flow, and canopy cover (Shah *et al*., 2017b). At high-elevations, an additional factor—the primary source of water input—plays an outsized role in dictating thermal variation downstream (Hotaling *et al*., 2017). High-elevation freshwaters are fed by four major hydrological sources: glaciers, snowfields, groundwater aquifers, and subterranean ice (Hotaling *et al*., 2019a, Tronstad *et al*., In press, Ward, 1994). Glaciers and subterranean ice (e.g., rock glaciers) promote near constant, extremely cold conditions (i.e., less than 3°C year-round) whereas snowmelt-and groundwater-fed streams are warmer and often more thermally variable (Hotaling *et al*., 2019a, Tronstad *et al*., In press). However, these general thermal “rules” apply only in close proximity to a primary source. Patterns can change dramatically downstream as flows are altered (e.g., pooling into a high-elevation pond) and sources mix (e.g., a warmer groundwater-fed stream flows into a glacier-fed stream). Resident aquatic taxa therefore experience vastly variable thermal conditions both within and across their life stages. With extensive thermal variation over small geographic scales and abundant, putatively cold-adapted resident invertebrates, high-elevation waters provide an ideal, natural model for testing hypotheses of physiological limits in a framework relevant to global change predictions.

In this study, we investigated gene expression as a function of tolerance to heat stress for stonefly nymphs collected from high-elevation streams in the northern Rocky Mountains. We focused on three taxa—*Lednia tetonica, Lednia tumana*, and *Zapada* sp.—all of which have habitat distributions closely aligned with cold, meltwater stream conditions. *Lednia tumana* was recently listed under the U.S. Endangered Species Act due to climate-induced habitat loss (U.S. Fish & Wildlife Service, 2019). To test tolerance to heat stress at the organism level, we measured the critical thermal maximum (CT_MAX_), a widely used metric for comparing thermal tolerance among animals (Healy *et al*., 2018). We specifically addressed three overarching questions: (1) Does natural thermal variation in stream temperature predict mountain stonefly CT_MAX_? (2) Do high-elevation stoneflies mount cellular stress responses when subjected to heat stress? And, if so, which genes are involved? (3) Is there a link between habitat conditions, organismal limits, and underlying gene expression? Following Shah *et al*. (2017b), we expected nymphs from streams with higher maximum temperatures to have correspondingly higher values of CT_MAX_. We also expected to observe a signal of cellular stress with genes typical of heat stress responses (e.g., HSPs) upregulated. Finally, we expected nymphs that naturally experience higher temperatures to exhibit a correspondingly muted cellular stress response. Collectively, our study sheds new light on thermal stress in high-elevation stream insects and contributes new perspective to a pressing challenge for the field: clarifying whether species living in cold headwaters are as sensitive to warming temperatures as their extreme distributions suggest.

## Materials and Methods

### Specimen collection

During the summer of 2018 (29 July-6 August), we collected late-instar stonefly nymphs representing at least three species (*Lednia tetonica, Lednia tumana*, and *Zapada* sp.; Family Nemouridae) from six streams in Glacier National Park (GNP), Montana, and Grand Teton National Park and the surrounding region (GRTE), Wyoming, USA (Figure 1; Tables 1, S1). We selected a later summer timepoint because it represents the warmest stream temperatures nymphs experience before emerging in August. Also, given the acclimation capacity of CT_MAX_ in temperate aquatic insects (Shah *et al*., 2017a), we measured CT_MAX_ during this period because it is also when we expected CT_MAX_ to be highest. Specimens were collected by turning over rocks and gently transferring nymphs to a small tray filled with streamwater. Nymphs were brought to the laboratory as quickly as possible in 1 L Whirl-Pak bags (Nasco) filled with streamwater surrounded by snow or ice. Species were identified based on morphological variation following previous studies (e.g., Giersch *et al*., 2017). Unlike *Lednia*, multiple *Zapada* species can be present in the same stream and previous genetic data has indicated the potential for cryptic diversity in the group (Hotaling *et al*., 2019b). Therefore, we cannot exclude the possibility of more than one species of *Zapada* in the Wind Cave population and thus only identified *Zapada* to genus (Table 1).

**Table 1.**
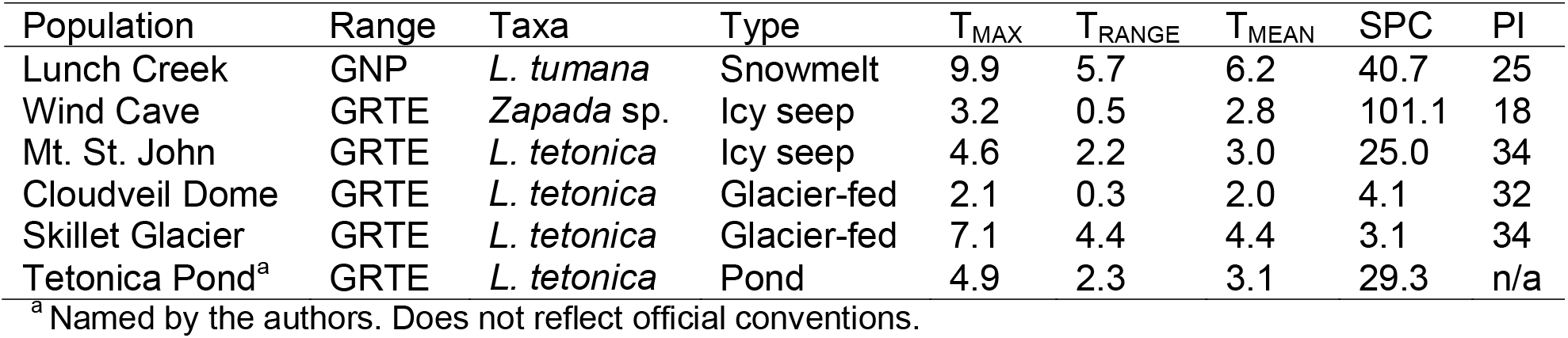
Environmental variation, mountain range, and habitat types included in this study. GNP: Glacier National Park, Montana. GRTE: Teton Range, Wyoming. T_MAX_: the maximum temperature observed, T_RANGE_: the difference between the maximum and minimum temperatures observed, and T_MEAN_: the mean temperature observed. All temperature data are in degrees Celsius. SPC: specific conductivity *μ*S cm^-1^), PI: Pfankuch Index, a measure of stream channel stability (higher values correspond to a less stable streambed). Temperatures were measured on a representative day in late July 2019 for all sites except Lunch Creek (data from late July 2014). See Table S1 for specific dates of temperature data collection.

**Figure 1.**
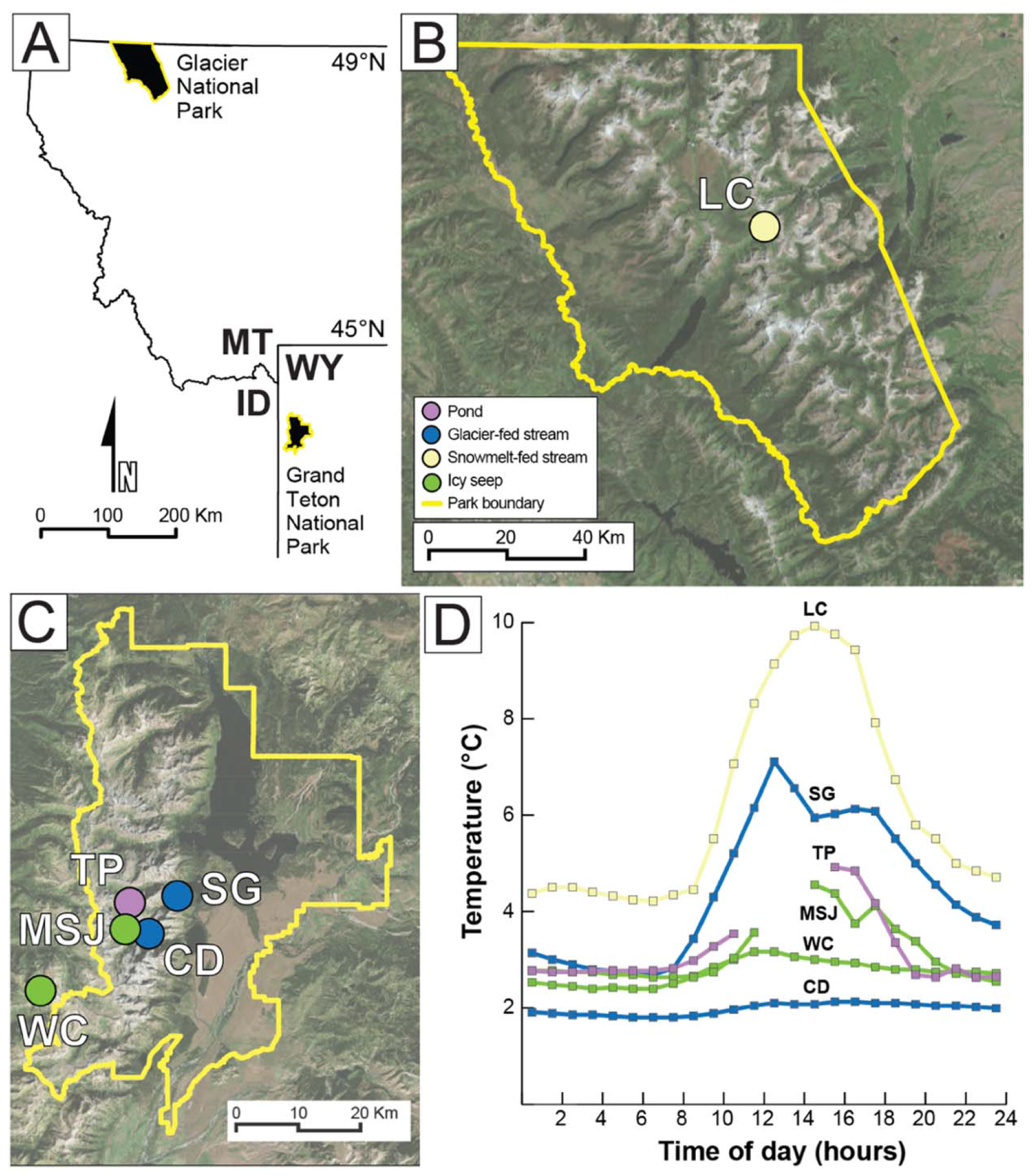
(A) The region of the Rocky Mountains where this study was conducted including (B) Glacier National Park, MT and (C) Grand Teton National Park, WY and the surrounding region. (D) A thermograph of hourly temperatures for each study site in late July. Site acronyms (top to bottom): Lunch Creek (LC), Skillet Glacier (SG), Tetonica Pond (TP), Mt. St. John (MSJ), Wind Cave (WC), and Cloudveil Dome (CD). A complete 24-hour thermograph is not shown for MSJ and TP because only 21 and 19 hours of continuous data were recorded for those sites, respectively. More extensive thermal data are provided in Figure S1.

### Environmental data and aquatic habitat classifications

For each study stream, we measured temperature by placing *in situ* HOBO loggers (Temperature Pro v2, Onset Computer Corporation) that recorded temperature hourly. Lengths of logger deployments ranged from less than 24 hours (Mt. St. John, Tetonica Pond) to several days (Cloudveil Dome) or a full year (Lunch Creek, Skillet Glacier, Wind Cave). Using these data, we constructed a one-day thermograph for each site based on a representative day in late July (exact dates provided in Table S1) and estimated the highest (T_MAX_), range (T_RANGE_), and mean (T_MEAN_) temperatures for that day. For two sites with more than one year of temperature data (Wind Cave: 2016, 2019; Lunch Creek: 2012, 2013, 2014), we compared multiple complete thermographs for July to ensure that our results were not biased by an unusual year- or day-specific pattern (Figure S1). We also collected two additional environmental variables to inform our habitat classifications (see below): specific conductivity (SPC), measured with a YSI Professional ProPlus multiparameter probe which was calibrated at the trailhead before each sampling trip, and stream channel stability, calculated via a modified version of the Pfankuch Index (PI), a standard metric for assessing channel stability in mountain systems that integrates five key physical characteristics of the stream into a single value (Peckarsky *et al*., 2014).

We classified sites into habitat types following previous studies (Giersch *et al*., 2017, Hotaling *et al*., 2019a, Tronstad *et al*., In press). Briefly, we incorporated a site’s primary hydrological source, environmental variation, and geomorphology, to group them into one of four habitat types: streams fed by a surface glacier (“glacier-fed”), a perennial snowfield (“snowmelt-fed”), emanating from subterranean ice (e.g., rock glaciers, “icy seep”), or slow-flowing, alpine ponds (“pond”). We categorized a stream as glacier-fed if it had a named glacier upstream and an extremely unstable streambed (PI > 30). Any other streams fed by perennial surface ice and snow were categorized as snowmelt-fed. We classified streams as icy seeps if we observed evidence of a subterranean ice source (e.g., lobes of a rock glacier), they were extremely cold (e.g., T_MAX_ < 5°C), and had high conductivity (SPC > 50; Hotaling *et al*., 2019a). Ponds were identified by their low-angle profile and the presence of standing water.

### Measuring critical thermal maxima (CT_MAX_)

Nymphs were brought into the laboratory as quickly as possible (typically less than 12 hours after collection) and transferred to holding chambers in 150-quart coolers filled with water from a nearby stream (Pacific Creek: 43.9036°, −110.5892°). We used aquarium chilling units (1/10 HP, Coralife) to maintain the holding baths at ~3°C (Figure S2). Each holding chamber contained 12 nymphs in a ~2 L plastic container immersed in the bath such that both water and nymphs were isolated from the rest of the system. We included plastic mesh in each chamber to give nymphs substrate to cling to. We maintained high levels of water flow and dissolved oxygen by air stone bubbling in each chamber. Nymphs had no access to food during the holding period to ensure they were tested in a fasting state (i.e., after available food had been digested, absorbed, and cleared from the digestive tract). All nymphs were held in these conditions for at least ~12 hours before testing (Table 2).

**Table 2.** Morphological and physiological data included in this study. Holding: time (hours) that specimens were held at 3°C with no access to food before testing. *N*: sample size for each population. Mean body lengths were used as a proxy for mass and are reported in millimeters with standard errors. RNAseq: sample sizes for RNA sequencing for treatment (T; CT_MAX_) and control (C; held at 3°C) specimens. Mean CT_MAX_ is given in degrees Celsius.

| Population | Taxon | Length | Holding | *N* | Mean CT<sub>MAX</sub> | RNAseq (*N*) |
| --- | --- | --- | --- | --- | --- | --- |
| Lunch Creek | *L. tumana* | 4.9 ± 0.5 | 72 | 24 | 28.7 | 3T / 3C |
| Wind Cave | *Zapada* sp. | 4.4 ± 0.6 | 48 | 23 | 25.9 | -- |
| Mt. St. John | *L. tetonica* | 5.6 ± 0.7 | 12 | 24 | 26.6 | 3T / 3C |
| Cloudveil Dome | *L. tetonica* | 4.5 ± 0.5 | 12 | 23 | 26.1 | -- |
| Skillet Glacier | *L. tetonica* | 5.6 ± 0.4 | 12 | 17 | 28.6 | -- |
| Tetonica Pond | *L. tetonica* | 4.6 ± 0.6 | 12 | 23 | 28.6 | 3T / 3C |

We measured CT_MAX_, a non-lethal temperature at which nymph locomotor function becomes disorganized. We placed up to 12 nymphs into mesh chambers (one individual per chamber) in a water bath held at 3°C. Because warming is the most obvious effect of climate change in high-mountain streams, we chose to only vary temperature, while maintaining natural flow and oxygenation with air pumps. Four thermo-electric cooling (TEC) plates attached to a temperature controller were used to increase temperature at ~0.25°C per minute. We recorded CT_MAX_ when an individual nymph could no longer right itself after being turned onto its back (Videos S1-S2). After a nymph reached its CT_MAX_, we immediately transferred it to an 8°C bath for recovery and assessed survival by monitoring nymphs until they resumed normal movement. Nymphs were later preserved in ~95% ethanol. We measured body length to the nearest ¼ mm using a dissecting microscope and a millimeter grid attached to the base of the microscope. A subset of nymphs were flash frozen at either their CT_MAX_ or holding temperature for RNA sequencing (RNAseq).

For CT_MAX_, all statistical analyses were conducted in R v3.4.0 (R Core Team, 2013). We focused our main analysis on *L. tetonica* because we had data for multiple populations. We first analyzed the effect of body size (length) and acclimation period on CT_MAX_ with linear models. Because body size was not a significant predictor of CT_MAX_ (see Results), our final linear models assessing the effect of maximum stream temperature (T_MAX_) on CT_MAX_ included only T_MAX_ as the predictor variable. We also assessed the effect of T_MAX_ on CT_MAX_ in a second, broader analysis which included the two populations of *L. tumana* and *Zapada* sp. While valuable, this ‘all-species’ analysis cannot be used to determine if CT_MAX_ varies among species or if the variation we observed is due to population-level differences because we lacked replicates for *L. tumana* and *Zapada* sp.

### RNA sequencing

During the thermal tolerance experiment, a subset of individuals from three populations and both *Lednia* species (Lunch Creek, *L. tumana*; Mt. St. John and Tetonica Pond, *L. tetonica*; Figure 1A, Table 2) were sampled for RNAseq. Nymphs at their CT_MAX_ (treatment) and others that remained at the holding temperature (control) were flash frozen in liquid nitrogen. We sampled three treatment and three control nymphs for each population (*N* = 18 total; Table 2). Samples were stored in liquid nitrogen until they were transferred to a −80° freezer. We extracted total RNA from entire nymphs following the NucleoSpin RNA (Macherey-Nagel Inc.) protocol. For extraction, specimens were re-flash frozen with liquid nitrogen in a 1.5 mL microcentrifuge tube and ground into a fine powder with a sterilized pestle. We quantified RNA with a Qubit 2.0 fluorometer (Thermo Fisher Scientific) and assessed RNA extraction quality via fragment analysis with an ABI 3730 DNA Analyzer (Thermo Fisher Scientific).

We prepared RNAseq libraries from 1 μg of total RNA with the NEBNext Poly(A) mRNA Magnetic Isolation Module (NEB) according to the manufacturer protocol. We targeted a 300-450 basepair (bp) fragment size distribution. For cDNA amplification, fifteen PCR cycles were used for all libraries. Presence of a PCR product was visually assessed using an eGel (Thermo Fisher Scientific). Final libraries were quantified with a Qubit 2.0 fluorometer and further assessed for quality, amount of cDNA, and fragment size distribution using a 2100 BioAnalyzer with the High Sensitivity DNA Analysis kit (Agilent). Libraries were then pooled in equal nanomolar concentrations and sequenced on one lane of an Illumina HiSeq4000 with 100 bp paired-end chemistry by the Roy J. Carver Biotechnology Center at the University of Illinois Urbana-Champaign.

### Gene expression analyses and protein annotation

We assessed raw sequence data quality with fastQC v0.11.4 (Andrews, 2010) and visualized a combined output for all libraries with MultiQC v1.5 (Ewels *et al*., 2016). Next, we trimmed reads in three successive rounds, all with Trim Galore! v0.4.1 (Krueger, 2015) and default settings except as noted. First, we removed adapter sequences (--illumina --stringency 6). Next, we trimmed for quality and poly-A tails (--quality 20 --stringency 6 --adapter A{30} -- adapter2 A{30}). We then trimmed for poly-T tails and discarded reads that had become too short (--stringency 6 --length 50 --adapter T{30} --adapter2 T{30}). We assessed the quality of the trimmed reads with fastQC v0.11.4. We randomly subsampled one library (Library 3; Control, Mt. St. John) to 80% of its original amount because its sequencing depth was much higher than the rest of the data set. For this, we used the reformat function of BBTools v37.80 (Bushnell, 2014). We removed one library (Library 9; Control, Mt. St. John) from all downstream analyses as it had just 2.6 million reads, far fewer than any other library (see Results).

We mapped reads to the *L. tumana* reference genome (GenBank #QKMV00000000.1) with the mitochondrial genome (GenBank #MH374046; Hotaling *et al*., 2019c) appended to it. We used HiSat2 v2.1.0 (Pertea *et al*., 2015) with default settings, first building an index of the reference with the hisat2-build command. To ensure no bias was introduced by differential mapping rates between *L. tumana* and *L. tetonica* samples to the *L. tumana* reference genome, we compared the mean mapping rates for both species with an unpaired *t*-test. Because HiSat2 outputs unsorted SAM files, we converted the output to sorted BAM files with samtools v1.7 (Li *et al*., 2009).

We generated a gene count matrix for each library with StringTie v1.3.5 (Pertea *et al*., 2015). We first ran StringTie with the default settings to assemble alignments into potential transcripts without a reference annotation (-G) because none is available for *L. tumana*. Next, we used the --merge utility to combine library-specific sets of transcripts into a merged, putatively non-redundant set of isoforms. This tool outputs a merged Gene Transfer Format (GTF) file. We then re-ran StringTie using the merged GTF (-G) and the flags −B and −e to enable the output of Ballgown GTF files for the global set of transcripts shared by all samples. Next, we ran the prepDE.py script, also part of the StringTie package, to generate counts matrices for all genes and transcripts identified in the previous steps.

We performed differential expression analyses using edgeR v3.26.8 (Robinson *et al*., 2010) in R version 3.5.2 (R Core Team, 2013). We filtered our data set by requiring transcripts to have more than five total reads and to be present in at least two samples. To visually compare expression variation across groups of interest (i.e., treatments, species, and populations), we used the plotPCA function. After filtering, we identified structure in global gene expression that could not be explained by sample preparation, library size, species, population, or treatment (Figure S3). We removed this unwanted variation with RUVseq v1.18.0 (Risso *et al*., 2014). Specifically, we used the *“in silico* empirical” functionality of RUVg where a set of the least differentially expressed genes (DEGs) are identified and used as controls to globally normalize variation in the data set. We used the default trimmed mean of M-values (TMM) method to normalize the data and calculate effective library sizes (Figure S4). Dispersions were estimated using a generalized linear model and a Cox-Reid profile-adjusted likelihood (McCarthy *et al*., 2012). We identified DEGs with quasi-likelihood F-tests (Lun *et al*., 2016) which were run using contrasts. We performed DEG identification across three levels of comparison: (1) Within-populations between treatment (collected at their CT_MAX_) and control (held at 3°C) specimens. (2) Between treatment and control for *L. tetonica* specimens only (Mt. St. John and Tetonica Pond). (3) Between treatment and control for all specimens. A false discovery rate (FDR) ≤ 0.05 was used to identify DEGs.

To annotate our data set, we extracted the longest isoform for each gene using the CGAT toolkit and the ‘gtf2gtf’ function (Sims *et al*., 2014). We then extracted genes from the file containing the longest isoforms with gffread v.0.9.9 (Trapnell *et al*., 2012). We performed a blastx search for each gene (E-value: 0.001) against the manually curated and reviewed Swiss-Prot database (Boeckmann *et al*., 2003; accessed 1 March 2019). Using the results of our blastx search, we annotated genes, retrieved gene ontology (GO) terms, and mapped GO terms using Blast2GO v5.2 (Conesa *et al*., 2005). We annotated DEGs with the top BLAST hit per transcript. For DEGs without a match in the Swiss-Prot database, we performed two additional searches using online tools: (1) a batch search against the RFAM database (Kalvari *et al*., 2017; http://rfam.org/search) and (2) a manual blast search (E-value: 0.001) against the automatically annotated and unreviewed TrEMBL database (Boeckmann *et al*., 2003; http://uniprot.org/blast/). In Blast2GO v5.2, we performed GO term enrichment analyses using the results of our blastx/Swiss-Prot annotations on two test sets with one-tailed Fisher’s Exact Tests and FDR ≤ 0.05 after correcting for multiple tests: (1) upregulated genes for *L. tetonica* only and (2) downregulated genes for *L. tetonica* only. We did not perform GO term enrichment analysis for *L. tumana* because no DEGs were identified for the representative population we examined (Lunch Creek; see Results). We also did not perform GO term enrichment on the overall *Lednia* data set because of redundancy with the *L. tetonica* analysis (i.e., roughly two-thirds of the same individuals would be included). For enrichment analyses, the complete set of transcripts with BLAST hits were used as the reference set.

To test if stoneflies from more thermally variable environments have muted cellular responses to stress, we identified all genes annotated as heat shock proteins based on BLAST hit descriptions. Next, we sorted these genes by their overall expression [log_2_ counts per million (logCPM)] and filtered them to a final set using two criteria: (1) We only included genes expressed at moderate to high levels (≥ 4 logCPM) and (2) only retained the most expressed hit (highest mean logCPM) for each unique gene. We did this to prevent any potential bias due to one gene being represented by multiple hits (see Results). Next, we calculated the mean difference in logCPM between treatment and control nymphs for each gene and population. Because the data were not normally distributed (*P*, Shapiro-Wilk < 0.001), we compared the distributions of mean differences for each population using a Kruskal-Wallis rank sum test followed by a Dunn test for multiple comparisons. All scripts and commands used in this study are available on GitHub (https://github.com/scotthotaling/Lednia_RNAseq).

## Results

### Environmental data and species collection

According to the environment criteria described above, we identified one snowmelt-fed stream (Lunch Creek: GNP), two icy seeps (Wind Cave, Mt. St. John; GRTE), two glacier-fed streams (Cloudveil Dome, Skillet Glacier; GRTE), and one alpine pond (Tetonica Pond; GRTE; Table 1). We collected *L. tumana* from Lunch Creek, *Zapada* sp. from Wind Cave, and *L. tetonica* from the other four sites (Figure 1, Table 1). Lunch Creek was the warmest (T_MEAN_ = 6.2°C; T_MAX_ = 9.9°C) and most thermally variable site (T_RANGE_ = 5.7°C; Table 1). Cloudveil Dome (T_MAX_ = 2.1°C) and Wind Cave (T_MAX_ = 3.2°C) were the coldest and least variable sites (T_RANGE_ ≤ 0.5°C; Table 1). Icy seeps were the coldest and least thermally variable habitat type overall (T_MAX_, icy seeps = 3.9°; T_RANGE_, icy seeps = 1.4°C). For the two sites with two or more years of available temperature data (2 years, Wind Cave; 3 years, Lunch Creek), thermal differences across years were negligible (Figure S1).

**Figure 2:**
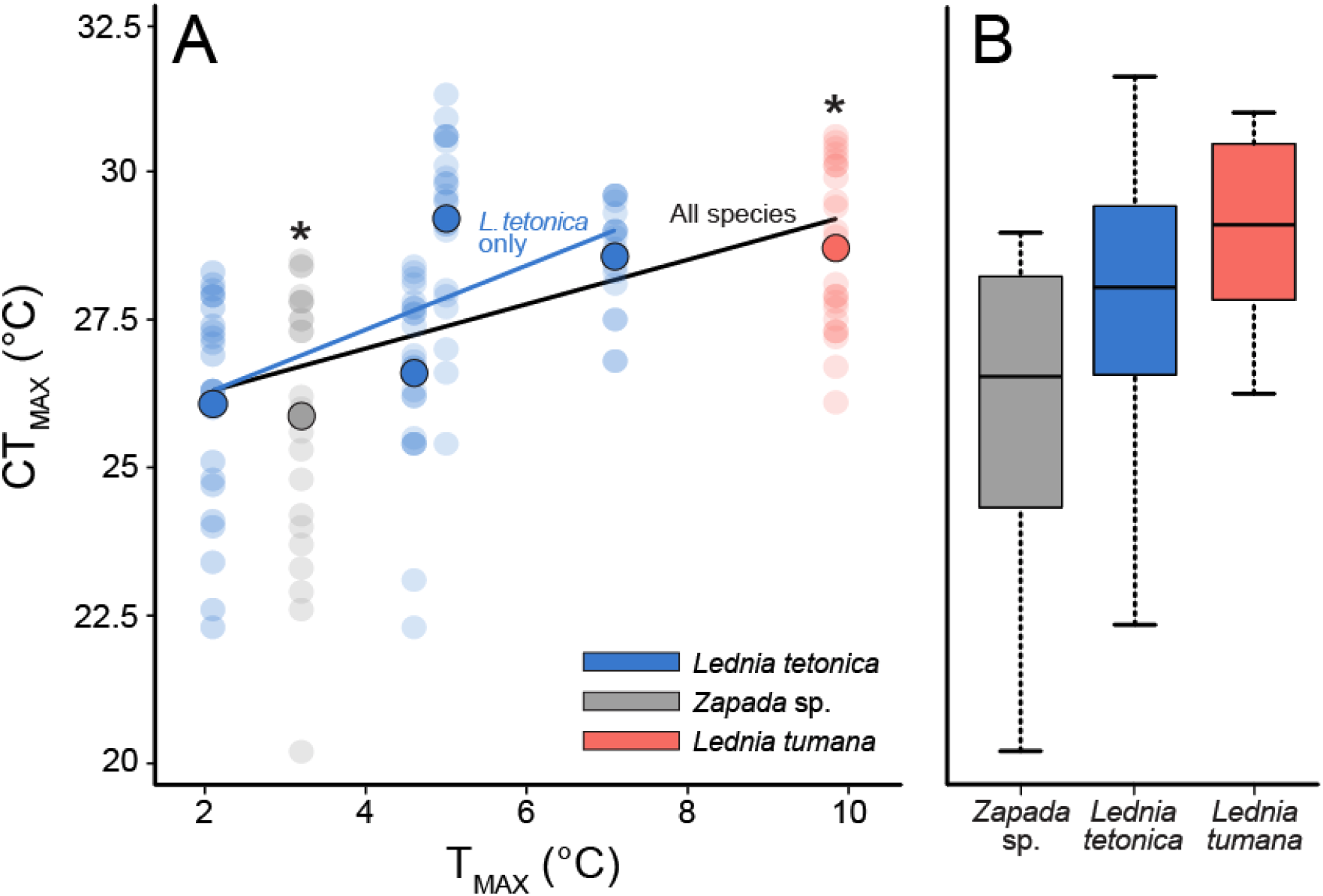
(A) The effect of maximum stream temperature (T_MAX_) on critical thermal maximum (CT_MAX_) for each nymph (smaller, lighter circles) with means for each population also shown (darker, outlined circles). Asterisks mark the species with only a single population sample (*Zapada* sp. and *L. tumana*). Trendlines indicate significant relationships between T_MAX_ and CT_MAX_ for two separate linear models for *L. tetonica* only (blue line) and for all species (black line). In both analyses, stoneflies from colder streams had lower CT_MAX_ values than those from warmer streams. (B) Box plots showing variation in CT_MAX_ across species. Black horizontal lines in each box indicate the median with lower and upper bounds of the box representing the lower and upper quartiles of the data, respectively. Whiskers show the maximum and minimum values.

### Thermal physiology

We confirmed that all nymphs survived the CT_MAX_ treatment (except for those that were immediately flash frozen for RNAseq and could not be assessed). Body size had no effect on CT_MAX_ in the *L. tetonica* (*P* = 0.58) nor all-species analysis (*P* = 0.28; Figure S5). We therefore did not include body size as a covariate in our statistical models. We also found no effect of acclimation period on CT_MAX_ (*P* = 0.41). We found differences in CT_MAX_ among populations of *L. tetonica* (Figure 2A). Stoneflies inhabiting colder sites exhibited lower CT_MAX_ values compared to those from warmer sites (*F*_1,85_ = 26.108, *P* < 0.001). We observed the lowest CT_MAX_ for *L. tetonica* in Cloudveil Dome (T_MAX_ = 2.1°C; CT_MAX_ = 26.1°C), and the highest for *L. tetonica* in Tetonica Pond (T_MAX_ = 5°C; CT_MAX_ = 29.2°C). We also found a significant positive relationship between T_MAX_ and CT_MAX_ in the all-species analysis which included *L. tumana* and *Zapada* sp. (*F*_1,132_ = 39.054, *P* < 0.001). Although we could not statistically test differences in CT_MAX_ among species due to a lack of replicate *L. tumana* and *Zapada* sp. populations, our results indicate that CT_MAX_ may be highest for *L. tumana* (Figure 2B). However, this finding may simply be reflective of the only *L. tumana* population sampled also being from Lunch Creek, the warmest site included in this study.

### RNA sequencing and annotation

We generated 368.8 million read pairs for 18 libraries with a mean per sample of 20.6 million ± 1.9 million (min. = 2.6 million, max. = 39.2 million). After filtering, subsampling of the library with the most reads, and dropping the library with the fewest reads, we retained 354.1 million read pairs. On average, 85.2% of reads mapped to the *L. tumana* reference genome with *L. tumana* libraries mapping at a slightly higher rate (mean 89.0% ± 0.5%; min. = 87.8%, max. = 91.0%) than *L. tetonica* (mean = 83.2% ± 0.6%; min. = 81.0%, max. = 84.5%; *P, t*-test < 0.0001). However, this difference in mapping rate did not extend to a difference in total reads mapped (mean, *L. tumana* = 19.2 million, mean *L. tetonica* = 21.7 million; *P, t*-test = 0.42). Raw reads for this study are deposited on the NCBI SRA under BioProject PRJNA587097.

### Differential expression

After filtering and processing of the data set, our gene counts matrix contained 52,954 unique entries. We observed global differences in gene expression between *L. tumana* and *L. tetonica* (Figure 3). When *L. tumana* and *L. tetonica* were combined (*“Lednia”*), 80 genes were differentially expressed: 65 upregulated and 15 downregulated in the treatment (CT_MAX_) versus control group (FDR ≤ 0.05). When only *L. tetonica* populations were considered (*“Tetonica”*), 71 genes were differentially expressed: 60 upregulated, 11 downregulated. Thirty-four DEGs were shared between groups (32 upregulated, two downregulated). When each population was considered alone, no DEGs were identified (including for Lunch Creek, the only *L. tumana* population). While we report results for the *Lednia* and *Tetonica* data sets above, we focus hereafter on *Tetonica* because it contains the most statistical power (two populations) with no potential for species-specific bias. Furthermore, due to the fragmented nature of the *L. tumana* genome (contig N50: 4.7 kilobases (kb); 74,445 contigs > 1 kb; Hotaling *et al*., 2019c), portions of the same gene were likely present on different contigs in the reference. When we assembled transcripts, this manifested as unique transcripts annotated to the same gene. Thus, in many instances (e.g., hexamerins, *HEXA*; Figures 4, S6), we recovered multiple independent hits to the same gene. While multiple hits may reflect biological reality (e.g., more than one copy of a gene in the genome perhaps reflecting a gene family expansion) we cannot draw such a conclusion. We specify how multiple hits to the same gene were handled where appropriate.

**Figure 3:**
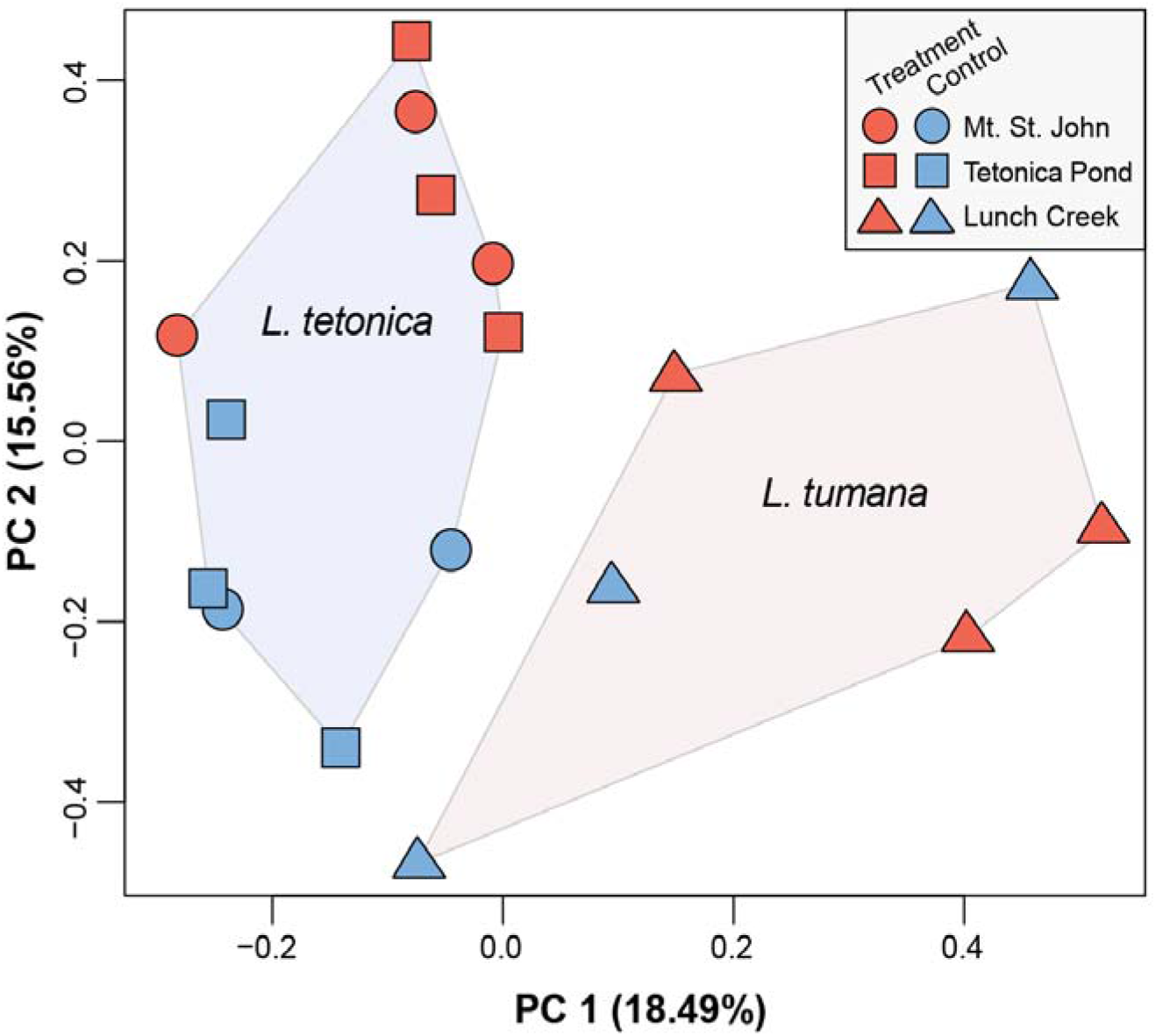
Global differences in gene expression for stonefly nymphs color-coded by treatment (red, CT_MAX_) or control (blue, held at 3°C) and grouped by species (colored polygons) and populations (shapes).

**Figure 4:**
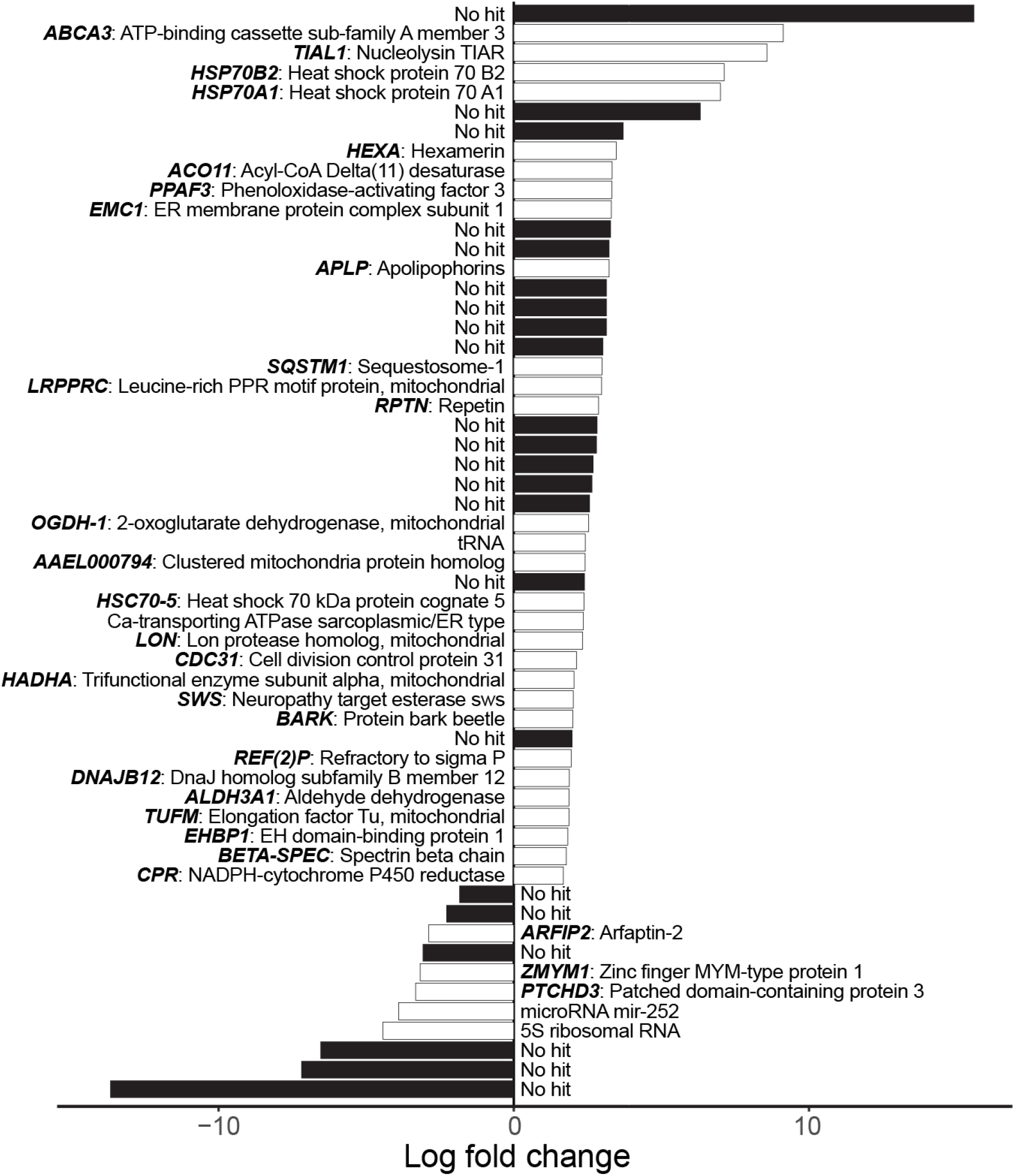
Log fold change of *Lednia tetonica* DEGs (white = BLAST annotated; black = no hit; FDR ≤ 0.05). For annotated genes, only the hits to the Swiss-Prot or RFAM databases with the lowest FDRs are included. The full version of this figure, including any instances of multiple hits to the same protein, is provided in Figure S6. Complete information for each annotation is provided in Table S2.

For *Tetonica*, 46 DEGs (64.8%) had BLAST hits to the high-quality, manually curated Swiss-Prot database, 32 of which were unique (Table S2). Of the remainder, three (4.2%) had hits to the RFAM database and nine (12.7%) had hits to the TrEMBL database. Because the TrEMBL protein database is unreviewed, we only refer to Swiss-Prot/RFAM annotations unless specifically noted. The most upregulated gene [MSTRG.32248; log_2_ fold change (logFC) = 15.6; FDR = 0.015] had no annotation to any database (Figure 4). However, the next four most-upregulated genes (logFC = 7-9.1; Figure 4) included *ABCA3*, which binds ATP, a nucleolysin (*TIAL1*), and two heat shock proteins *HSP70B2* and *HSP70A1*. The two heat shock proteins were also the most expressed DEGs (logCPM = 8.9 and 9.3, respectively) after three genes which were all annotated as hexamerins (*HEXA*; logCPM = 9.3-10). Fourteen DEGs had hits to the same apolipophorin gene, *APLP*, with relatively similar changes in expression (logFC, *APLP* = 2.1-3.8; Figure S6) and overall expression levels (logCPM, *APLP* = 2.2-6.9). The three most downregulated genes did not have BLAST hits to the Swiss-Prot or RFAM databases [logFC = −6.6 to −13.7; Figure 4], however two of them (MSTRG.1867 and MSTRG.3534) had TrEMBL hits though they were not informative in terms of predicted function (Table S2).

Forty-one GO terms were enriched in the upregulated *Tetonica* data set (Figure S7): 26 were classified as being part of a biological process ontology, three were cellular component related, and 11 were linked to molecular function. The top four most significantly enriched GO terms were all lipid-related, including their transport, binding, and localization. Eight of the enriched GO terms (19.5% overall) were associated with protein folding, and three were linked to chaperone proteins which are commonly associated with physiological stress (Beissinger & Buchner, 1998). In the same vein, one enriched GO term – “heat shock protein binding” (GO:0031072; FDR = 0.015) – clearly reflected a link to heat stress at the cellular level. No GO terms were enriched for downregulated *Tetonica* DEGs.

### Environmental variability and gene expression

Across all populations and species, 38 genes were annotated as heat shock proteins (HSPs). Of these, 12 unique genes were expressed at moderate to high levels (logCPM ≥ 4; Figure S8). We found no support for our hypothesis that stoneflies naturally experiencing higher (and more variable) temperatures exhibit muted cellular stress responses versus those inhabiting colder (and more thermally stable) streams (Figure 5; *P*, Dunn’s ≥ 0.66).

**Figure 5:**
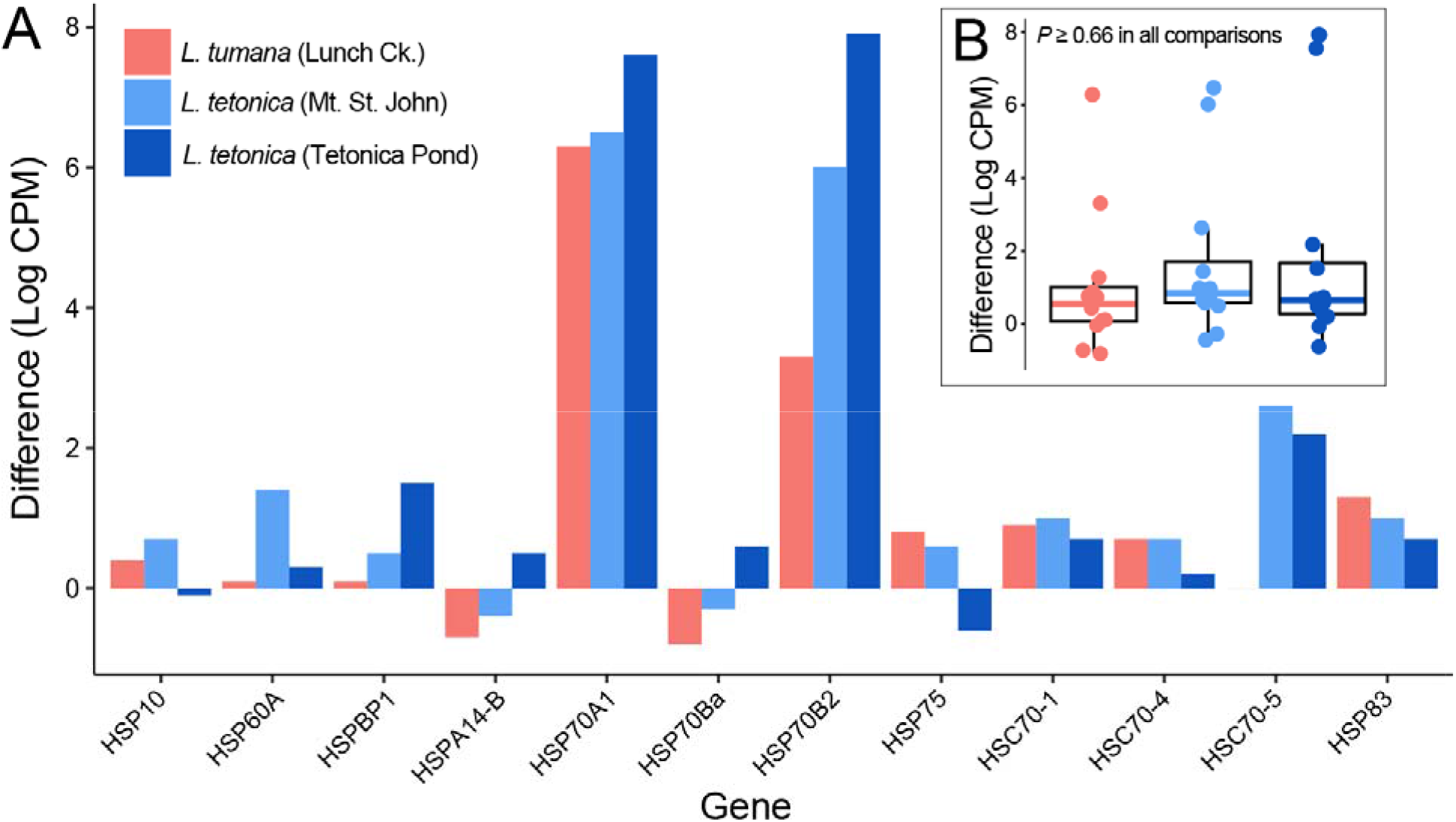
(A) Mean population-level differences in expression between treatment and control specimens for the 12 most highly expressed, unique HSPs annotated in this study. (B) Distributions of the values in (A) grouped by population. No significant differences were present (*P*, Dunn < 0.05).

## Discussion

As climate change proceeds, headwaters will be dramatically altered by the reduction or loss of meltwater from glaciers and perennial snowfields (Hotaling *et al*., 2017). However, the physiological limits of high-elevation aquatic insects, a group presumed to be acutely imperiled by climate change, remain largely unknown. In this study, we explored the thermal physiology of high-elevation stoneflies inhabiting the meltwater of rapidly fading glaciers and snowfields in the Rocky Mountains. Our focal species are representative of an entire community that may be at risk of climate-induced extirpation (Giersch *et al*., 2017, Hotaling *et al*., 2019a, Tronstad *et al*., In press), and included *L. tumana*, a species listed under the U.S. Endangered Species Act due to climate-induced habitat loss (U.S. Fish & Wildlife Service, 2019). We show that habitat thermal conditions, specifically maximum temperatures, predict upper thermal limits and that nymphs mount a cellular stress response when faced with heat stress. Contrary to our expectations, however, we saw no link between the scale of the stress response and natural conditions that nymphs experience. That is, stoneflies from warmer environments did not exhibit a muted cellular stress response across HSPs versus those from cooler streams. Our results shed new light on thermal tolerance of mountain stoneflies and complement recent cellular perspectives on aquatic insect thermal biology (Ebner *et al*., 2019, Gamboa *et al*., 2017). Broadly, our findings and those of others (e.g., Ebner *et al*., 2019, Muhlfeld *et al*., 2020, Shah *et al*., 2017b, Treanor *et al*., 2013), challenge the prevailing notion that aquatic insect larvae living in extremely cold mountain streams cannot survive warming. For *Lednia*, with the ability to tolerate short-term temperature spikes, we hypothesize that their headwater distributions may be a product of longer-term physiological mechanisms (e.g., the capacity to develop at near freezing temperatures) and biotic factors (e.g., species interactions at lower elevation).

### Thermal tolerance

In mountain systems, thermal tolerance is important to organismal distributions and can help explain the elevation limits of many terrestrial taxa (Andrews, 1998, Brattstrom, 1968, Feder & Lynch, 1982, Huey & Webster, 1976, Oyen *et al*., 2016). Whether thermal tolerance can also explain distributional limits of aquatic taxa is unknown (Polato *et al*., 2018). We show that species of high-elevation stoneflies in the Rocky Mountains, often described as cold stenotherms that are highly susceptible to warming (e.g., Giersch *et al*., 2017), can withstand relatively high short-term temperatures (see also Shah *et al*., 2017b). Although the utility of CT_MAX_ has been challenged due to its sensitivity to ramping rates, as well as acclimation and starting temperatures (Rezende *et al*., 2011, Terblanche *et al*., 2011), recent arguments in favor of its ecological relevance have also been made (Jørgensen *et al*., 2019), especially when used in a comparative framework. We contend that CT_MAX_ may be uniquely appropriate for mountain stream taxa. Indeed, alpine streams are rapidly warming (e.g., Niedrist & Füreder, 2020) and in our study system, swift increases in temperature (over a few hours) are common (e.g., Lunch Creek, Figure 1D). With summer streamflows predicted to be reduced under climate change (Huss & Hock, 2018), baseline temperatures and intraday temperature spikes will both increase as meltwater volume declines and its buffering capacity is lost.

In addition to temperature spikes during certain seasons, average alpine stream temperatures are on the rise (Niedrist & Füreder, 2020). These higher, but sublethal, temperatures will likely have pervasive negative impacts on high elevation aquatic insects (Shah *et al*., 2019). For example, long-term thermal tests of *L. tumana* development suggest that mortality during emergence greatly increases around 15 °C (A.A.S. & S.H., unpublished data), highlighting how subtle thermal effects on ecological timescales may limit the persistence of *L. tumana* and similar species. Long-term temperature shifts will likely have complex effects on larval energy budgets by differentially altering rates, as well as targets, of energy expenditure (e.g., resource allocation between somatic growth, maintenance, and reproduction) and energy income from feeding.

Simultaneous increases in temperature and reductions in flow may also elevate heat sensitivity in mountain stoneflies by exacerbating a mismatch between oxygen supply and demand. For ectotherms, increasing temperature typically results in increased metabolic rates and greater demand for oxygen (Verberk *et al*., 2016a). In aquatic habitats, this demand may not be met with adequate oxygen supply, eventually leading to decreased organismal fitness (Pörtner & Knust, 2007). Evidence for this phenomenon, however, is mixed (Verberk *et al*., 2016b). With so little known of aquatic insect thermal physiology, it is imperative for future research to address effects of sublethal temperatures and oxygen limitation on thermal tolerance, especially in high-elevation aquatic insects that may encounter a lethal combination of increased temperatures and decreased oxygen from low flows (Jacobsen, 2020).

We observed variation in CT_MAX_ among populations of *L. tetonica*, suggesting that local thermal regime may be more important to thermal tolerance than regional thermal regime, and echoing the findings of other recent studies (Gutiérrez-Pesquera *et al*., 2016, Shah *et al*., 2017b). This effect of local conditions on thermal tolerance might outweigh differences due to evolutionary history because all species (e.g., *Lednia tetonica* and *Zapada* sp.) from cooler streams had lower CT_MAX_ than those from warmer streams (Figure 2). Although we cannot determine if thermal variation among populations represents evolved differences, all specimens were held in a common thermal regime for at least 12 hours to limit the effects of previous thermal conditions on CT_MAX_ estimates. Regardless of the mechanism, the high-elevation stonefly nymphs included in this study appear poised to cope with short-term warming in streams, although some populations are likely to be more resilient than others (e.g., those experiencing higher present-day maximum temperatures).

Given that we focused on larvae in our study, we cannot discern whether other life stages (e.g., eggs or adults) differ in their thermal tolerance. However, we focused on nymphs for three reasons. First, the larval stage is the key developmental period for aquatic insects when the majority of growth and other fitness-related processes (e.g., egg production) occur, and recent modeling suggests that impacts of climate change on species can be greatly *underestimated* when the larval stage is overlooked (Levy *et al*., 2015). Second, aquatic insect larvae typically do most of their growing during summer months, when the threat of heat stress is greatest. And, third, egg hatching success in the laboratory was recently shown to be consistently high for mountain stream insects, regardless of rearing temperature, including one treatment (12°C) that exceeded the highest temperatures the focal midges naturally experienced (Schütz & Füreder, 2019). Still, future experiments that link traits (e.g., thermal tolerance) to cellular processes across aquatic insect life cycles will greatly improve our understanding of how sensitivity at key life stages may influence long-term viability of populations.

### Gene expression

High-elevation stoneflies residing in cold meltwater-fed streams exhibited a cellular stress response when faced with temperatures at their CT_MAX_. The bulk of this response was comprised of upregulated genes and included well-known stress response genes (e.g., HSPs; Lindquist & Craig, 1988), lesser known but potentially stress-related genes in insects (e.g., APLP, Dassati *et al*., 2014), and many DEGs that could not be annotated (Figure 4). Three HSPs (*HSP70B2, HSP70A1, HSC70-5*) were upregulated in nymphs experiencing thermal stress. With well-established roles as cellular protectants, preventing protein denaturation, binding aberrant proteins, and many other stress-induced measures, the upregulation of HSPs was unsurprising (King & MacRae, 2015). However, given the seemingly psychrophilic lifestyle of *Lednia*, where individuals develop at temperatures near 0°C, we expected to see widespread upregulation of HSPs in treatment nymphs. This was not the case. Rather, *Lednia* appeared to constitutively express many HSPs even at low temperature (Figure S8). This suggests that, contrary to the prevailing view, exposure to low temperatures may actually stress *Lednia* (see additional discussion below). Similar patterns of constitutive HSP expression have been observed in other cold-tolerant species. For instance, larval caddisflies (Ebner *et al*., 2019), polar fish (Buckley *et al*., 2004), and Antarctic grass (Reyes *et al*., 2003) constitutively express many HSPs, presumably to chaperone proteins at low temperature. The potential for *Lednia* to be stressed by cold temperatures is further supported by the inability of *L. tumana* nymphs to survive contact with ice (Hotaling *et al*., In review).

We also observed upregulation of genes with lipid-related functions. Lipids, particularly those in membranes, are extremely sensitive to changes in temperature (Hazel, 1995), and because of their role in key biological processes (e.g., solute diffusion), are important for thermal acclimation and adaptation (Muir *et al*., 2016, Pernet *et al*., 2007). Many ectotherms remodel their membrane lipids under thermal stress to maintain fluidity, typically through increases in saturated fatty acids at higher temperatures (Muir *et al*., 2016). The process of maintaining membrane fluidity in the face of changing temperatures has been termed homeoviscous adaptation (HVA; Sinensky, 1974). The degree of saturation in cuticular lipids of stoneflies varies across life stages, and was speculated to be related to thermal tolerance, particularly as it relates to the aquatic versus terrestrial environment (Armold *et al*., 1969). To our knowledge, the presence of HVA has not been explicitly tested for in any aquatic insect. However, before upregulation of lipid-related genes can be presumed to underlie HVA or similar functional changes in high-elevation stoneflies, alternate explanations about the role of lipids in development must be considered (see below).

While heat stress is presumed to drive the expression patterns we observed, aquatic insects accelerate their development and emerge earlier at warmer temperatures (Nebeker, 1971, Rempel & Carter, 1987), sometimes even during CT_MAX_ experiments (A.A.S., personal observation). Thus, some expression changes may be the result of developmental shifts rather than thermal stress directly. When exposed to long-term temperatures above those they naturally experience (e.g., ≥ 15°C for ~1 month), *L. tumana* nymphs rapidly develop compared to those held at colder temperatures. However, rapidly emerging adults can get stuck and die while shedding their cuticle (S.H. and A.A.S., unpublished data). Some of our results appear more reflective of this developmental shift than heat stress directly. For instance, it has been suggested that *ABCA3* is upregulated during insect wing development (Broehan *et al*., 2013). In our study, high temperatures induced upregulation of *ABCA3*, perhaps indicating accelerated wing development in preparation for adult emergence. Lipid content in aquatic insects also varies seasonally and tends to peak before metamorphosis, an energetically demanding activity (Cavaletto & Gardner, 1999). In the caddisfly, *Clisoronia magnifica*, roughly 80% of the lipid reserves needed for metamorphosis were used during the last instar (Cargill *et al*., 1985), which was the same developmental timepoint of the stoneflies included in this study.

The upregulation of *HEXA* raises similar, albeit more complex, questions. Stoneflies possess two types of hexameric proteins in their hemolymph: hemocyanin (*HCD*), an oxygen-carrying protein, and hexamerins, multi-functional proteins that likely evolved from hemocyanin (Amore *et al*., 2011, Hagner-Holler *et al*., 2007). We saw some evidence for the upregulation of *HCD* in heat-stressed stoneflies (Figure S9), perhaps reflecting the physiological challenge of extracting the necessary oxygen from warmer water. However, while hexamerins likely evolved from *HCD*, their function shifted to storage proteins after they lost the ability to bind oxygen (Burmester, 2015, Markl & Winter, 1989). Present-day hexamerins primarily act as sources of amino acids during non-feeding periods (e.g., emergence, Haunerland, 1996) but may also play a role in cuticle formation (Burmester, 2015, Hagner-Holler *et al*., 2007), a key stage in aquatic insect emergence. Thus, the upregulation of *HEXA* may be another cellular indicator of accelerated emergence to escape injurious conditions.

### Mountain stream insects as cold stenotherms: reconsidering a historical paradigm

Aquatic insects living in chronically cold habitats have long been assumed to be cold-adapted and therefore intolerant of warming (e.g., Giersch *et al*., 2017, Jacobsen *et al*., 2012). This assumption has rarely, if ever, been supported by direct measurements. A potential mismatch between theory and data is particularly important for imperiled species of conservation concern. *Lednia tumana* is federally endangered under the U.S. Endangered Species Act due to loss of cold, meltwater habitat (U.S. Fish & Wildlife Service, 2019). As glaciers disappear around the world (Huss & Hock, 2018), the demise of *Lednia* and similar species (e.g., *Zapada* sp.) is presumed to be merely a matter of time (Giersch *et al*., 2017). While this may be true, alternative hypotheses or threats beyond temperature, at least in the short term, should be considered. Chief among these is the question of realized niche breadth. Factors limiting niche breadth are diverse and may not be directly linked to temperature (e.g., interspecific competition or food availability, Connell, 1961, Roughgarden, 1974), although thermal sensitivity can certainly play a major role (Gilchrist, 1995). While terrestrial habitats exhibit a wide array of thermal variation, potentially allowing more thermal space for species with similar ecologies to exist in sympatry, the buffering capacity of flowing water may reduce the diversity of thermal niches in streams across similar spatial extents (Shah *et al*., 2020). Thus, if *Lednia* exhibit high short-term thermal tolerance, exceeding temperatures they naturally experience, and cellular signatures of stress even at low temperatures (e.g., constitutive expression of HSPs at 3°C), then we hypothesize that the distribution of *Lednia* and similar species reflects not a requirement for cold conditions but simply a greater tolerance for them versus other species. Rather than being an extreme thermal specialist, *Lednia* may have evolved a wide thermal niche allowing it to colonize environments free of limiting biological factors. Our hypothesis aligns with previous experimental evidence highlighting the potential for biotic factors beyond temperature to alter alpine stream ecosystems (Khamis *et al*., 2015) and large-scale ecological data showing the persistence of meltwater-associated biodiversity after deglaciation (Muhlfeld *et al*., 2020).

When considering climate change impacts on mountain stream biodiversity, it is important to distinguish between a species imperiled by rising temperatures, biotic factors, or seemingly by a combination of the two (e.g., Durance & Ormerod, 2010). At present, the prevailing theory is that a warmer water community will shift uphill and displace coldwater taxa as glaciers and perennial snowfields are lost (Hotaling *et al*., 2017). This theory assumes that coldwater species (e.g., *Lednia*) will not be able to tolerate warmer conditions and will be extirpated while lower elevation species simultaneously track their preferred thermal conditions upstream. However, if existing headwater communities can tolerate warmer conditions and their lower limits are set by other factors (e.g., competition), then climate change risks for mountain stream communities may be far less generalizable than currently assumed. For instance, if competition at lower elevations limits *Lednia* distributions then warming temperatures do not guarantee simplistic, binary outcomes of predicted presence or absence. Rather, the future of *Lednia* and similar taxa may depend upon whether their competitors shift uphill (rather than tolerate warmer conditions *in situ*), how resources may change, and additional factors that are difficult to predict (see Shah *et al*., 2020).

## Conclusion

High-elevation stoneflies in the Rocky Mountains can tolerate higher temperatures in the short-term than those they experience in the wild. When challenged with heat stress, nymphs mount a cellular response that includes upregulation of classic stress response genes (e.g., HSPs) as well as genes that may be involved in developmental transitions from aquatic to terrestrial life stages. Aquatic insects, including *L. tumana*, develop more rapidly at stable warmer temperatures but also experience higher mortality during emergence (Nebeker, 1971; S.H. and A.A.S., unpublished data). Thus, the potential effects of sublethal warming on performance and other fitness-related traits warrant further investigation. However, in light of our results and similar studies (Ebner *et al*., 2019, Muhlfeld *et al*., 2020, Shah *et al*., 2017b), we challenge the premise that the distribution of mountain stream insects in cold, thermally stable habitats indicates specialized preferences for cold, or evolved physiologies that are only viable in the cold. Rather, the appearance of constitutive expression of many HSPs in *Lednia* as well as the inability of *L. tumana* nymphs to survive ice enclosure (Hotaling *et al*., In review) indicate their contemporary thermal regimes may actually be injurious. Ultimately, if imperiled species like *Lednia* are not directly threatened by warming temperatures in the near term, then there is clear reason for greater optimism about their future. However, explicit investigations of their development under warmer regimes, rather than simplistic, short-term exposures, are needed in concert with new understanding of how other abiotic factors (e.g., oxygen supply), biotic interactions, and resource availability shape their distributions.

## Acknowledgements

We thank the University of Wyoming-National Park Service (UW-NPS) Research Station for funding. S.H. and J.L.K. were supported by NSF awards #IOS-1557795 and #OPP-1906015. A.A.S. was supported by an NSF Postdoctoral Research Fellowship in Biology (DBI-1807694).

M.E.D. was supported by NSF awards #DEB-1457659, #OIA-1826834, and #EF-1921562. Harold Bergman, Winsor Lowe, Taylor Price, and Lydia Zeglin provided valuable logistic, laboratory, or field assistance. The UW-NPS Research Station provided laboratory space to perform the experiments and the Ghalambor Lab provided insect holding equipment. We performed computational analyses on the Kamiak High Performance Computing Cluster at Washington State University. We thank Christopher Kozakiewicz, Stephen Ormerod, Mark Smithson, and three anonymous reviewers for comments that improved the manuscript. Any use of trade, firm, or product names is for descriptive purposes only and does not imply endorsement by the U.S. Government.

## Author contributions

S.H. and A.A.S. conceived of the study. S.H., A.A.S., K.L.M., L.M.T., J.J.G., D.S.F., M.E.D., and J.L.K. collected the data. S.H. and A.A.S. analyzed the data and wrote the manuscript with input from K.L.M., H.A.W., M.E.D., and J.L.K. All authors read and approved the final version.

