## Supplementary Materials for "Mountain stoneflies may tolerate warming streams: evidence from organismal physiology and gene expression"

**Supplementary Tables:**

**Table S1.** GPS coordinates and elevations of study sites. Elevations are in meters. GNP: Glacier National Park, GRTE: Grand Teton National Park and surrounding mountains. Temperature data indicates the date(s) when T_MAX_, T_RANGE_, and T_MEAN_ in Table 1 were measured from a representative ~24-hour period. Temperature data for Mt. St. John and Tetonica Pond span two days because loggers were placed on July 30^th^ and retrieved the following day. Elevations are reported in meters.

| Population | Range | Temperature data | Latitude, Longitude | Elevation |
| --- | --- | --- | --- | --- |
| Lunch Creek | GNP | July 31, 2014 | 48.704, -113.703 | 2,090 |
| Wind Cave | GRTE | July 28, 2019 | 43.666, -110.961 | 2,692 |
| Mt. St. John | GRTE | July 30-31, 2019 | 43.791, -110.785 | 2,690 |
| Cloudveil Dome | GRTE | July 31, 2019 | 43.725, -110.796 | 2,927 |
| Skillet Glacier | GRTE | July 31, 2019 | 43.840, -110.755 | 2,690 |
| Tetonica Pond | GRTE | July 30-31, 2019 | 43.786, -110.800 | 2,883 |

**Table S2.** Annotations for all differentially expressed genes identified for *L. tetonica*. Grayed out entries denote hits to the same gene which were not included in Figure 4 (but are included in Figure S6). Since BLAST searches were made against both of the UniProtKB databases – Swiss-Prot (high-quality; manually annotated and reviewed) and TrEMBL (automatically annotated and not reviewed) – the accession ID is given with the database in parentheses (Swiss-Prot = SP; TrEMBL = T). Gene names reflect those of the UniProtKB database.

| Transcript ID | LogFC | LogCPM | *P* | FDR | Gene | Description | UniProtKB ID |
| --- | --- | --- | --- | --- | --- | --- | --- |
| MSTRG.32248.1 | 15.59 | 6.46 | 6.09E-06 | 0.0151 | n/a | None | n/a |
| MSTRG.18905.1 | 9.12 | 0.08 | 3.80E-05 | 0.0429 | ABCA3 | ATP-binding cassette sub-family A member 3 | Q99758 (SP) |
| MSTRG.11533.1 | 8.57 | 5.04 | 4.16E-05 | 0.0429 | TIAL1 | Nucleolysin TIAR | Q01085 (SP) |
| MSTRG.3952.1 | 7.13 | 8.87 | 9.77E-08 | 0.0013 | HSP70B2 | Heat shock protein 70 B2 | P41827 (SP) |
| MSTRG.2749.1 | 7.00 | 9.31 | 1.31E-08 | 0.0006 | HSP70A1 | Heat shock protein 70 A1 | P41825 (SP) |
| MSTRG.29029.1 | 6.32 | -0.34 | 1.32E-05 | 0.0252 | n/a | Acyltransferase | A0A2P8XQ87 (T) |
| MSTRG.15964.1 | 3.77 | 2.63 | 2.24E-05 | 0.0335 | n/a | Apolipophorins | Q9U943 (SP) |
| MSTRG.53594.1 | 3.71 | 2.34 | 2.29E-06 | 0.0087 | n/a | None | n/a |
| MSTRG.50157.1 | 3.69 | 3.06 | 4.89E-05 | 0.0429 | n/a | Apolipophorins | Q25490 (SP) |
| MSTRG.55919.1 | 3.50 | 4.01 | 1.91E-05 | 0.0311 | n/a | Apolipophorins | Q25490 (SP) |
| MSTRG.32216.1 | 3.47 | 9.75 | 3.11E-05 | 0.0388 | n/a | Hexamerin | Q17127 (SP) |
| MSTRG.17593.1 | 3.40 | 9.33 | 4.99E-05 | 0.0429 | n/a | Hexamerin | Q17127 (SP) |
| MSTRG.5174.1 | 3.33 | 5.61 | 1.42E-06 | 0.0087 | n/a | Acyl-CoA Delta(11) desaturase | Q8ISS3 (SP) |
| MSTRG.5025.1 | 3.33 | 2.97 | 4.73E-05 | 0.0429 | PPAF3 | Phenoloxidase-activating factor 3 | Q8I6K0 (SP) |
| MSTRG.3532.1 | 3.31 | 5.53 | 6.07E-08 | 0.0011 | EMC1 | ER membrane protein complex subunit 1 | Q5ZL00 (SP) |
| MSTRG.40754.1 | 3.29 | 3.21 | 3.01E-06 | 0.0098 | n/a | Apolipophorins | A0A067R667 (T) |
| MSTRG.42883.1 | 3.23 | 4.40 | 3.56E-07 | 0.0031 | n/a | None | n/a |
| MSTRG.51312.1 | 3.23 | 3.86 | 5.47E-07 | 0.0041 | n/a | Apolipophorins | Q9U943 (SP) |
| MSTRG.32016.1 | 3.23 | 5.91 | 2.17E-06 | 0.0087 | n/a | Apolipophorins | Q9U943 (SP) |
| MSTRG.44357.1 | 3.15 | 3.78 | 2.97E-05 | 0.0388 | n/a | Apolipophorins | Q9U943 (SP) |
| MSTRG.18854.1 | 3.15 | 2.68 | 4.95E-05 | 0.0429 | n/a | None | n/a |
| MSTRG.50833.1 | 3.14 | 3.78 | 2.88E-06 | 0.0098 | n/a | None | n/a |
| MSTRG.31276.1 | 3.14 | 0.67 | 5.06E-05 | 0.0429 | n/a | None | n/a |
| MSTRG.27467.1 | 3.02 | 4.44 | 1.59E-07 | 0.0017 | n/a | ETS domain-containing protein | A0A2J7R7V7 (T) |
| MSTRG.28428.1 | 3.00 | 5.66 | 3.45E-06 | 0.0100 | n/a | Apolipophorins | Q9U943 (SP) |
| MSTRG.38342.1 | 2.98 | 4.75 | 5.79E-05 | 0.0449 | SQSTM1 | Sequestosome-1 | O08623 (SP) |
| MSTRG.12688.2 | 2.97 | 6.16 | 1.62E-05 | 0.0272 | LRPPRC | Leucine-rich PPR motif-containing protein, mitochondrial | Q5SGE0 (SP) |
| MSTRG.18178.1 | 2.90 | 2.19 | 1.96E-05 | 0.0311 | n/a | Apolipophorins | Q9U943 (SP) |
| MSTRG.404.1 | 2.87 | 2.38 | 6.77E-05 | 0.0499 | RPTN | Repetin | Q6XPR3 (SP) |
| MSTRG.56303.1 | 2.87 | 2.46 | 2.90E-05 | 0.0388 | n/a | Apolipophorins | Q9U943 (SP) |
| MSTRG.28641.1 | 2.85 | 6.62 | 1.81E-06 | 0.0087 | n/a | Apolipophorins | Q9U943 (SP) |
| MSTRG.48990.1 | 2.82 | 4.07 | 2.34E-06 | 0.0087 | n/a | None | n/a |
| MSTRG.39006.1 | 2.81 | 5.64 | 1.97E-06 | 0.0087 | n/a | Peroxidase | A0A287J6B7 (T) |
| MSTRG.21296.1 | 2.70 | 6.89 | 1.81E-06 | 0.0087 | n/a | Apolipophorins | Q9U943 (SP) |
| MSTRG.47266.1 | 2.69 | 3.40 | 5.63E-05 | 0.0449 | n/a | None | n/a |
| MSTRG.8905.1 | 2.65 | 3.05 | 5.09E-05 | 0.0429 | n/a | ETS domain-containing protein | A0A2J7R7V7 (T) |
| MSTRG.27504.1 | 2.57 | 1.48 | 6.60E-05 | 0.0493 | n/a | Heat shock protein 75kDA, mitochondrial | A0A2J7PFP3 (T) |
| MSTRG.20968.1 | 2.53 | 4.31 | 4.02E-05 | 0.0429 | OGDH-1 | 2-oxoglutarate dehydrogenase, mitochondrial | O61199 (SP) |
| MSTRG.33031.1 | 2.42 | 6.72 | 7.02E-06 | 0.0167 | n/a | Apolipophorins | Q9U943 (SP) |
| MSTRG.45077.1 | 2.42 | 5.55 | 1.36E-05 | 0.0252 | n/a | tRNA (non-coding RNA) | n/a |
| MSTRG.5820.2 | 2.41 | 6.54 | 3.24E-06 | 0.0100 | AAEL000794 | Clustered mitochondria protein homolog | Q17N71 (SP) |
| MSTRG.18456.1 | 2.40 | 3.63 | 4.75E-05 | 0.0429 | n/a | Peptidase_M3 domain-containing protein | A0A2J7R330 (T) |
| MSTRG.5646.1 | 2.37 | 6.68 | 4.19E-05 | 0.0429 | HSC70-5 | Heat shock 70 kDa protein cognate 5 | P29845 (SP) |
| MSTRG.51801.3 | 2.37 | 10.04 | 3.43E-05 | 0.0416 | n/a | Hexamerin | Q17127 (SP) |
| MSTRG.6707.1 | 2.36 | 7.87 | 9.97E-06 | 0.0217 | SERCA | Ca-transporting ATPase sarcoplasmic/ER type | Q7PPA5 (SP) |
| MSTRG.8870.2 | 2.33 | 5.24 | 2.37E-05 | 0.0341 | LON | Lon protease homolog, mitochondrial | Q7KUT2 (SP) |
| MSTRG.37392.1 | 2.14 | 5.72 | 1.19E-05 | 0.0250 | n/a | Apolipophorins | Q9U943 (SP) |
| MSTRG.25311.1 | 2.12 | 3.19 | 2.99E-05 | 0.0388 | CDC31 | Cell division control protein 31 | P06704 (SP) |
| MSTRG.51499.1 | 2.10 | 5.33 | 4.56E-05 | 0.0429 | n/a | Apolipophorins | Q9U943 (SP) |
| MSTRG.9376.1 | 2.04 | 5.10 | 1.40E-05 | 0.0252 | HADHA | Trifunctional enzyme subunit alpha, mitochondrial | P40939 (SP) |
| MSTRG.23204.1 | 2.01 | 2.19 | 5.05E-05 | 0.0429 | SWS | Neuropathy target esterase sws | B4M709 (SP) |
| MSTRG.3402.1 | 2.00 | 3.89 | 5.09E-06 | 0.0133 | BARK | Protein bark beetle | M9NDE3 (SP) |
| MSTRG.26346.1 | 1.98 | 4.31 | 7.34E-06 | 0.0167 | n/a | None | n/a |
| MSTRG.19199.1 | 1.94 | 6.96 | 4.02E-05 | 0.0429 | REF(2)P | Refractory to sigma P | P14199 (SP) |
| MSTRG.28037.1 | 1.88 | 2.65 | 6.44E-05 | 0.0488 | DNAJB12 | DnaJ homolog subfamily B member 12 | Q9NXW2 (SP) |
| MSTRG.24681.1 | 1.87 | 4.36 | 4.40E-05 | 0.0429 | ALDH3A1 | Aldehyde dehydrogenase, dimeric NADP-preferring | P30838 (SP) |
| MSTRG.41001.1 | 1.86 | 3.69 | 5.76E-05 | 0.0449 | TUFM | Elongation factor Tu, mitochondrial | Q8BFR5 (SP) |
| MSTRG.19752.1 | 1.83 | 3.34 | 3.99E-05 | 0.0429 | EHBP1 | EH domain-binding protein 1 | Q69ZW3 (SP) |
| MSTRG.1982.1 | 1.78 | 3.18 | 2.18E-05 | 0.0335 | BETA-SPEC | Spectrin beta chain | Q00963 (SP) |
| MSTRG.20797.1 | 1.68 | 4.21 | 5.07E-05 | 0.0429 | CPR | NADPH-cytochrome P450 reductase | Q27597 (SP) |
| MSTRG.52236.1 | -1.84 | 4.05 | 5.84E-05 | 0.0449 | n/a | None | n/a |
| MSTRG.36087.1 | -2.28 | 2.77 | 5.22E-05 | 0.0433 | n/a | None | n/a |
| MSTRG.13112.2 | -2.88 | 3.54 | 4.88E-05 | 0.0429 | ARFIP2 | Arfaptin-2 | Q3ZCL5 (SP) |
| MSTRG.16003.1 | -3.08 | 1.92 | 2.41E-05 | 0.0341 | n/a | None | n/a |
| MSTRG.7951.1 | -3.18 | 4.69 | 5.33E-05 | 0.0435 | ZMYM1 | Zinc finger MYM-type protein 1 | Q5SVZ6 (SP) |
| MSTRG.35645.1 | -3.33 | 2.94 | 3.78E-05 | 0.0429 | PTCHD3 | Patched domain-containing protein 3 | Q0EEE2 (SP) |
| MSTRG.28832.1 | -3.91 | 1.25 | 3.08E-05 | 0.0388 | n/a | microRNA mir-252 (non-coding RNA) | n/a |
| MSTRG.55446.1 | -4.43 | 0.19 | 1.36E-05 | 0.0252 | n/a | 5S ribosomal RNA (ncRNA) | n/a |
| MSTRG.1867.1 | -6.55 | -1.57 | 1.50E-05 | 0.0262 | n/a | TPR_MLP1_2 domain-containing protein | A0A1Y1V8P0 (T) |
| MSTRG.31287.1 | -7.20 | -0.99 | 3.74E-06 | 0.0103 | n/a | None | n/a |
| MSTRG.3534.1 | -13.67 | 4.28 | 2.35E-08 | 0.0006 | n/a | ATP-grasp_3 domain-containing protein | A0A1Z4S904 (T) |

**Supplementary Figures:**

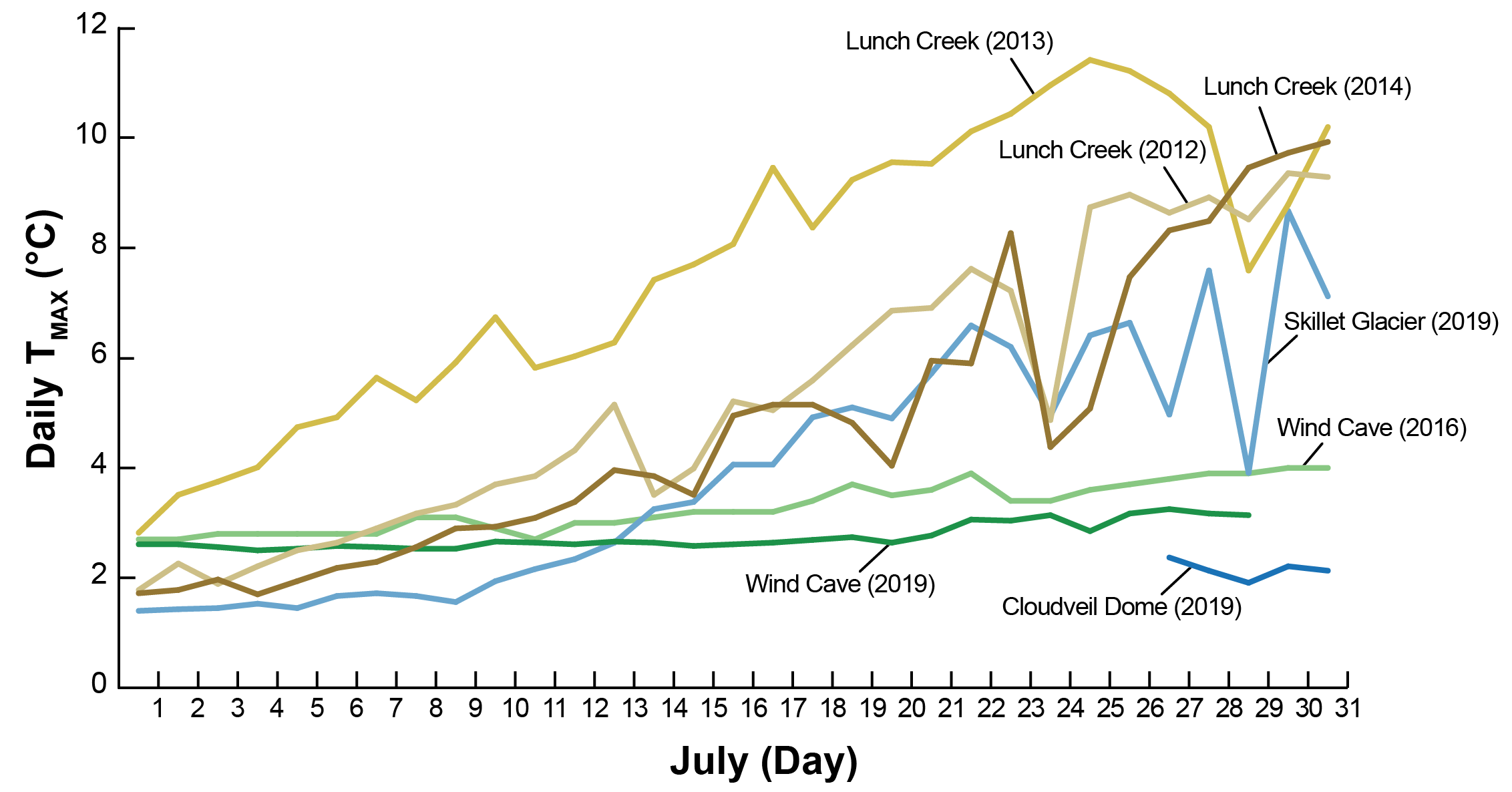

**Figure S1.** Maximum daily temperatures (T_MAX_) throughout July for study sites with three or more consecutive days of temperature data collection. More than one July of data are shown for Wind Cave (2016, 2019) and Lunch Creek (2012, 2013, 2014) highlighting that our focal data (a representative day in late July, see Table S1) were unlikely to be biased by a year- or day-specific pattern.

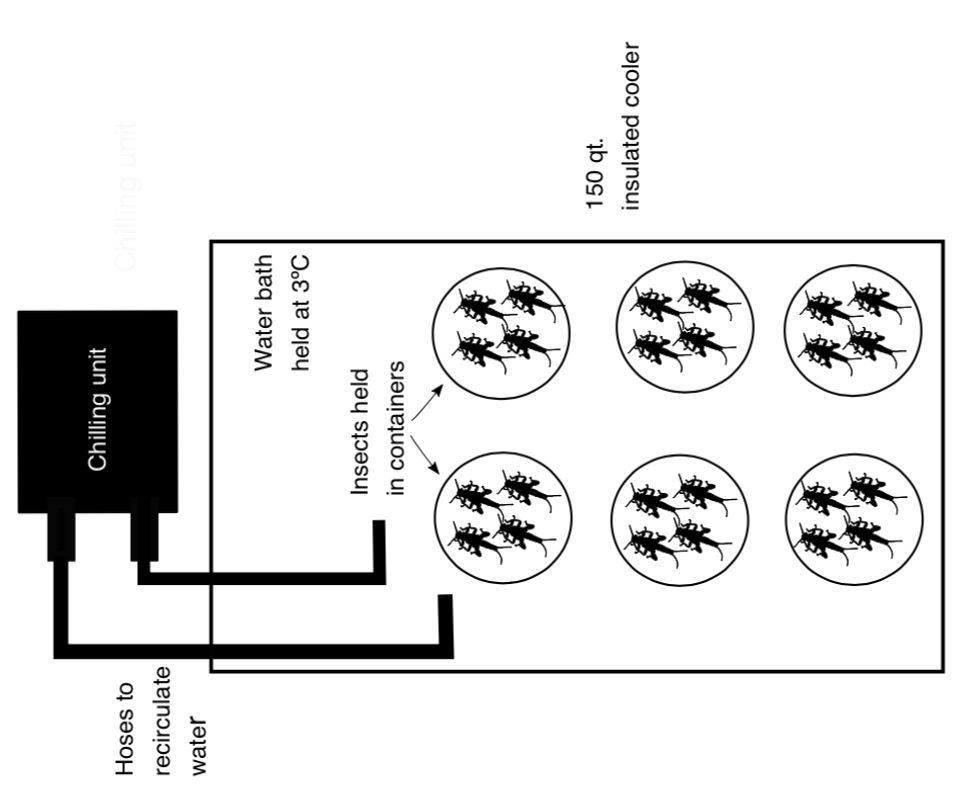

**Figure S2**. Schematic of our laboratory holding set up for wild-caught stoneflies. Nymphs were held at 3ºC without food for 12 hours. Note that due to logistical challenges, populations from Wind Cave and Lunch Creek were held for 48 and 72 hours, respectively.

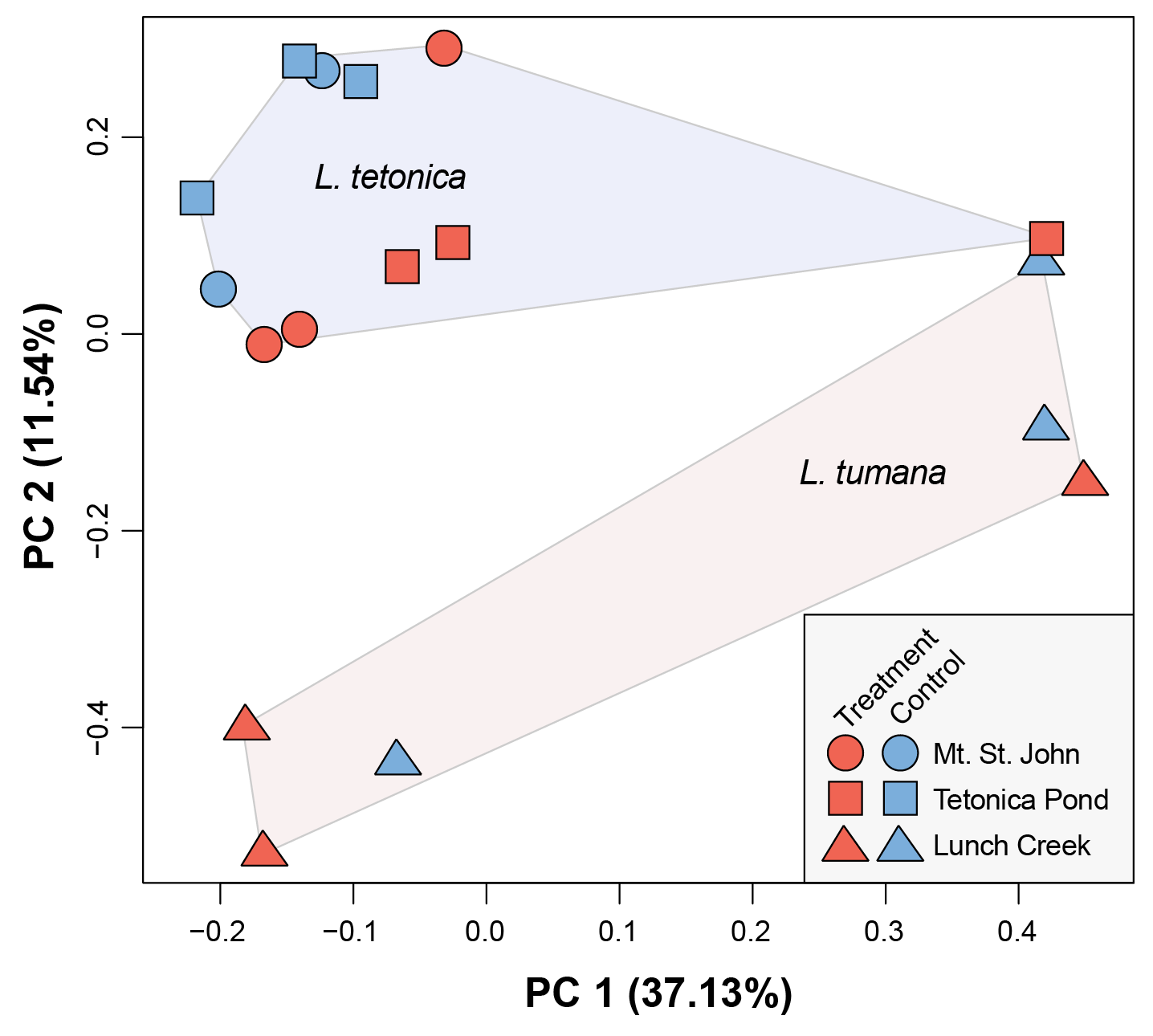

**Figure S3**. Global differences in gene expression before unexplained variation was removed with RUVseq.

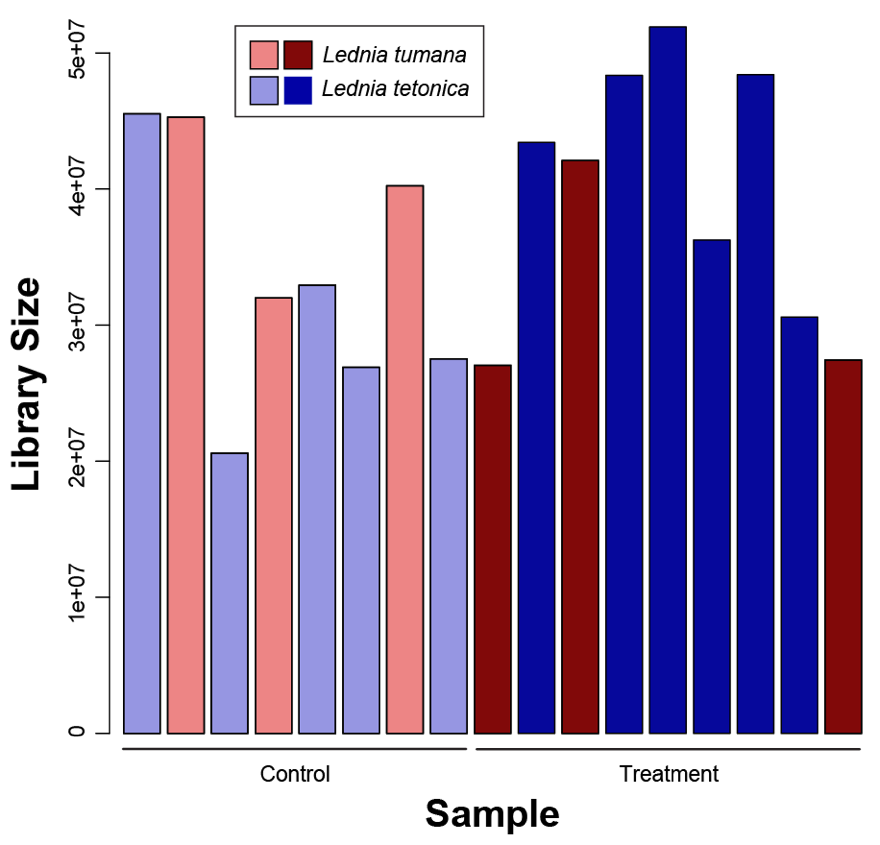

**Figure S4.** Effective library size for each sample as calculated with the trimmed mean of M-values method. The effective library size is equal to the product of the original library size multiplied by a scaling factor which minimizes the log-fold change between samples for the most genes.

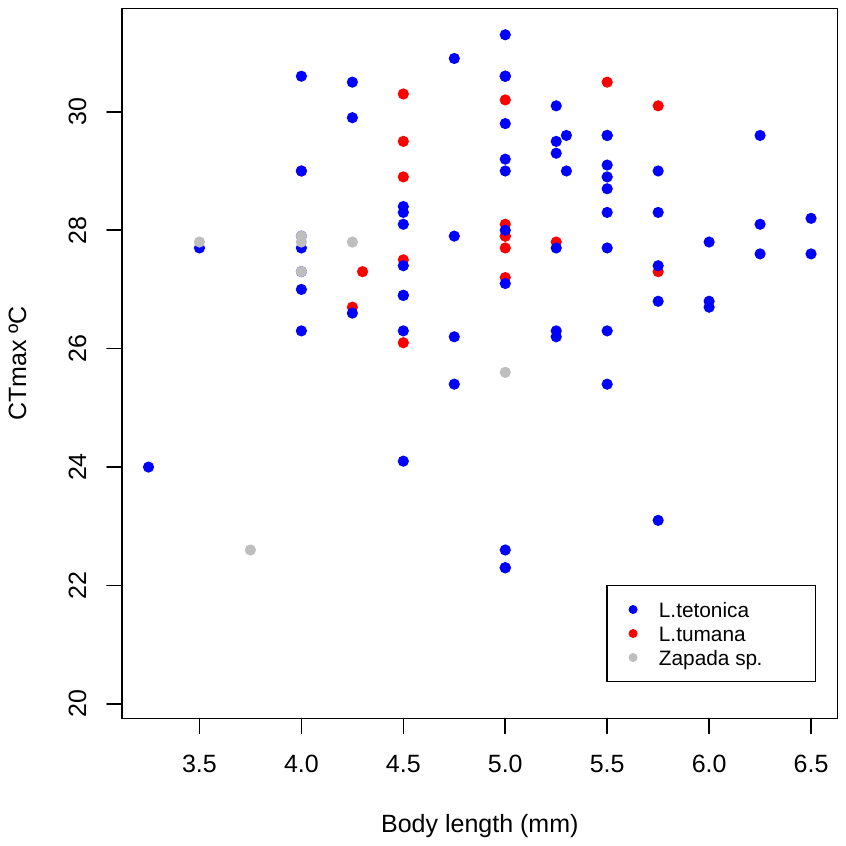

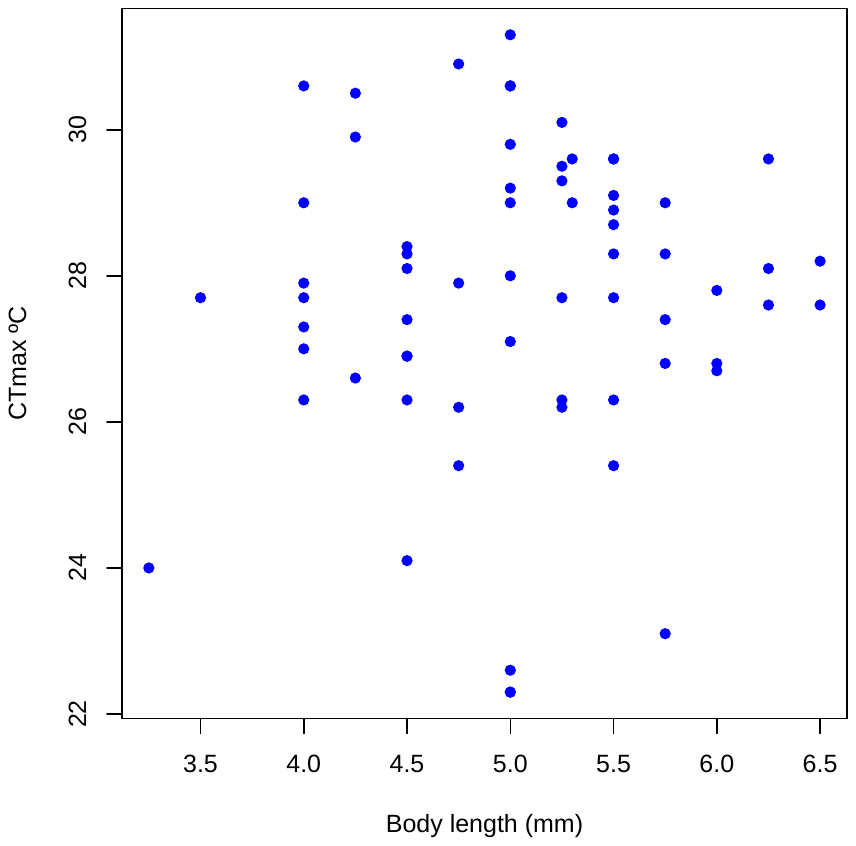

**B**

**A**

**Figure S5**. A regression of CT_MAX_ on body length for (A) *L. tetonica* only and (B) all species. Each circle represents one nymph. We did not find an effect of body length on CT_MAX_ for *L tetonica* (*F*_1,63_ = 0.315, *P* = 0.58) nor across all species (*F*_1,89_ = 1.202, *P* = 0.28). We therefore did not include body length as a parameter in any of our statistical models.

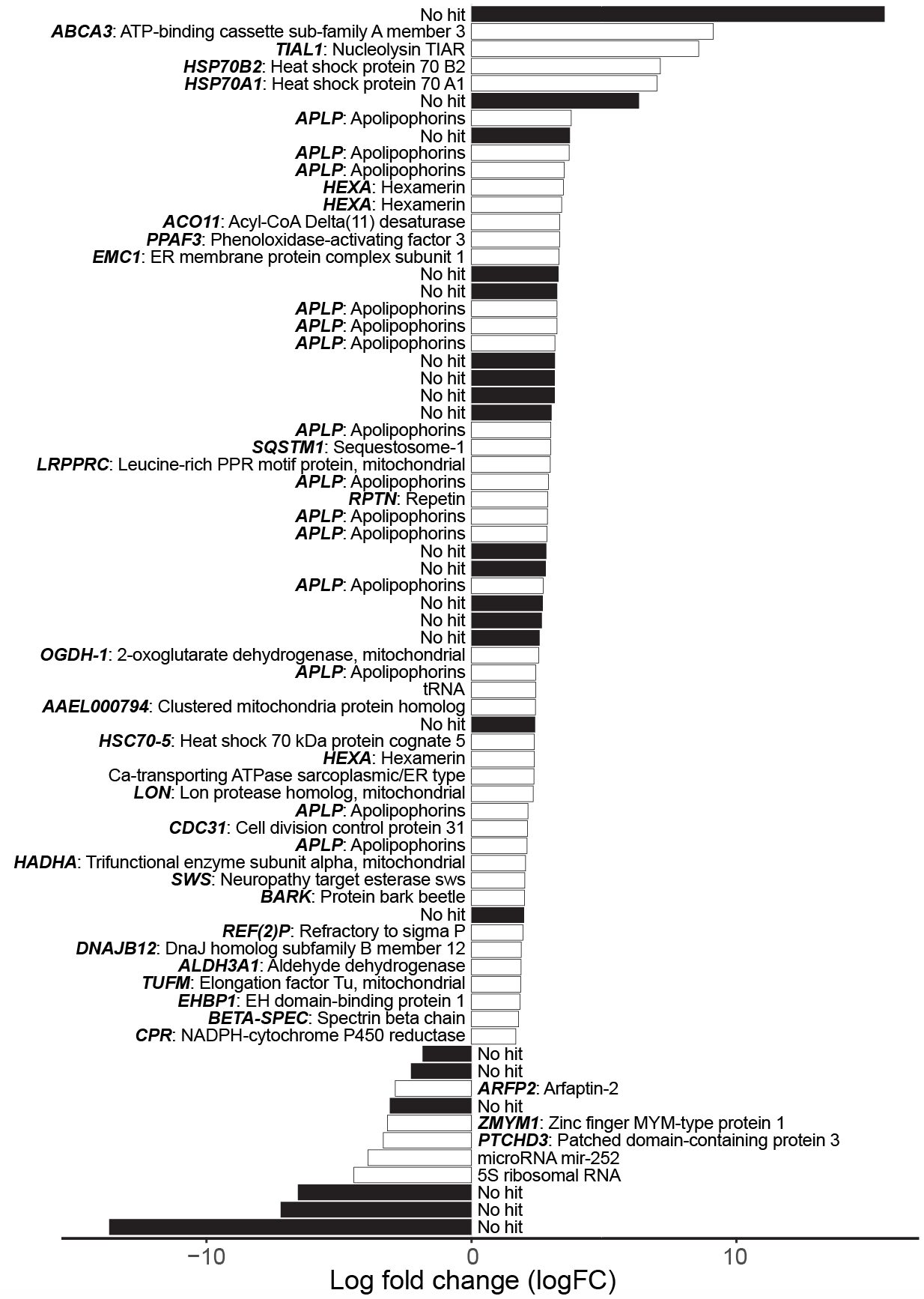

**Figure S6.** Log fold change of *Lednia tetonica* proteins (white = annotated; black = no hit) that were differentially expressed (FDR ≤ 0.05). Duplicate hits (e.g., hexamerins) are the result of either multiple copies in the *L. tumana* genome or errors due to fragmentation of the reference. Only hits to the Swiss-Prot or RFAM databases are included. The complete list of annotations is included in Table S2.

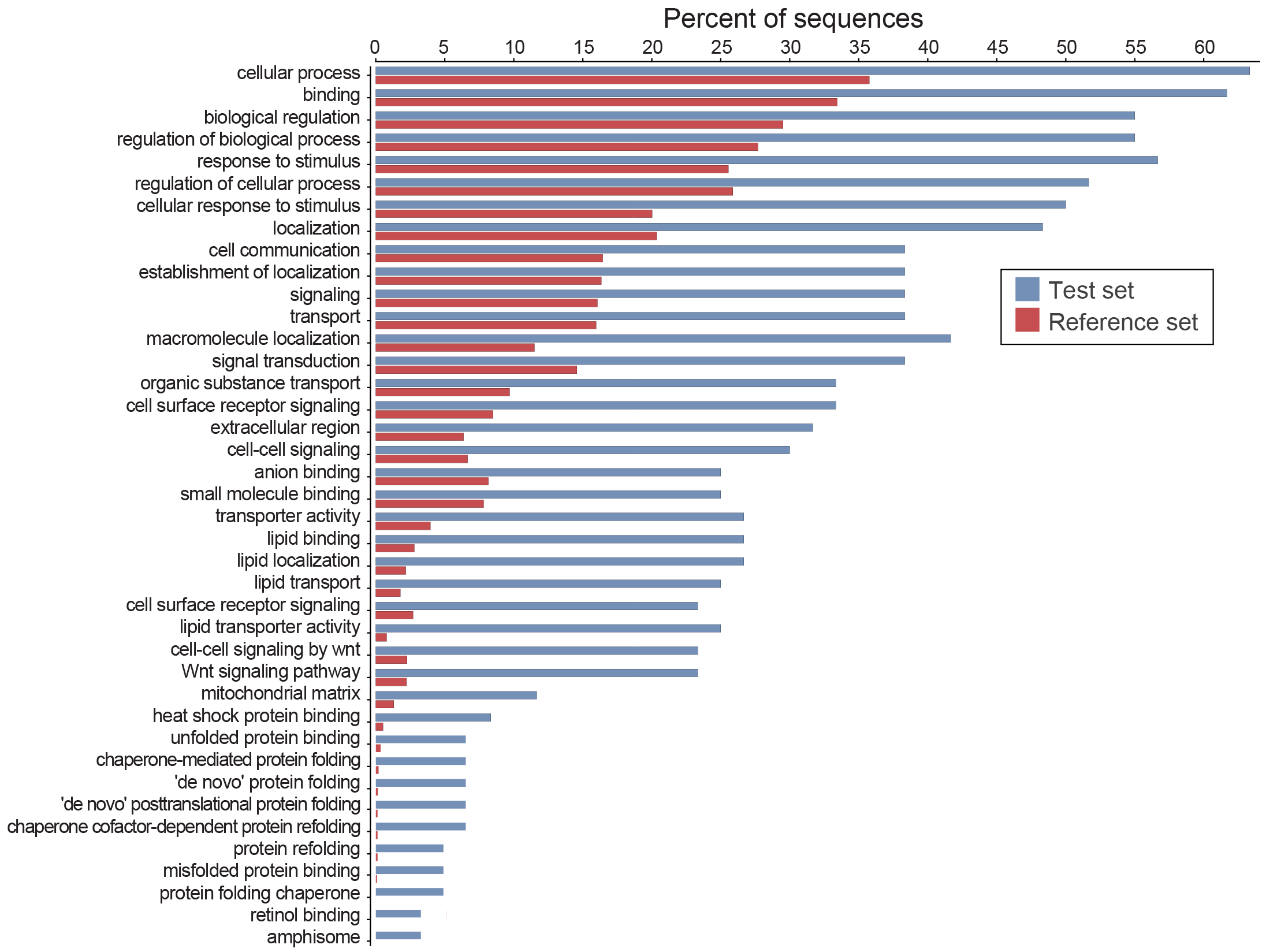

**Figure S7.** Significantly enriched gene ontology terms for *L. tetonica* differentially expressed genes (test set) compared to all genes in the data set (reference set).

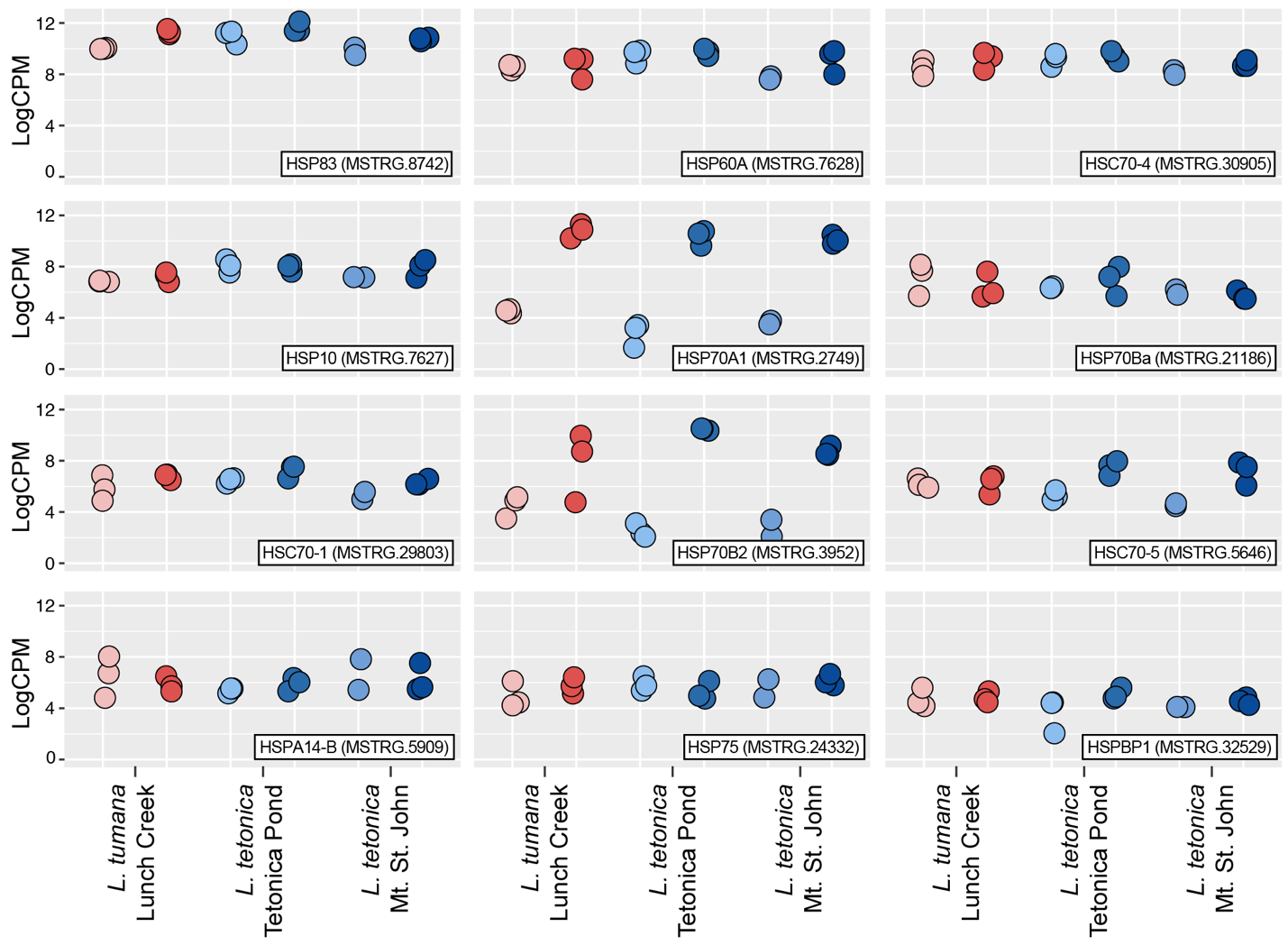

**Figure S8.** Gene expression [log_2_ counts per million (logCPM)] for 12 heat shock proteins (gene annotation in bottom right corner of each plot) between control (lighter colors) and treatment (darker colors) nymphs. Most heat shock proteins are­ constitutively expressed in *Lednia.*

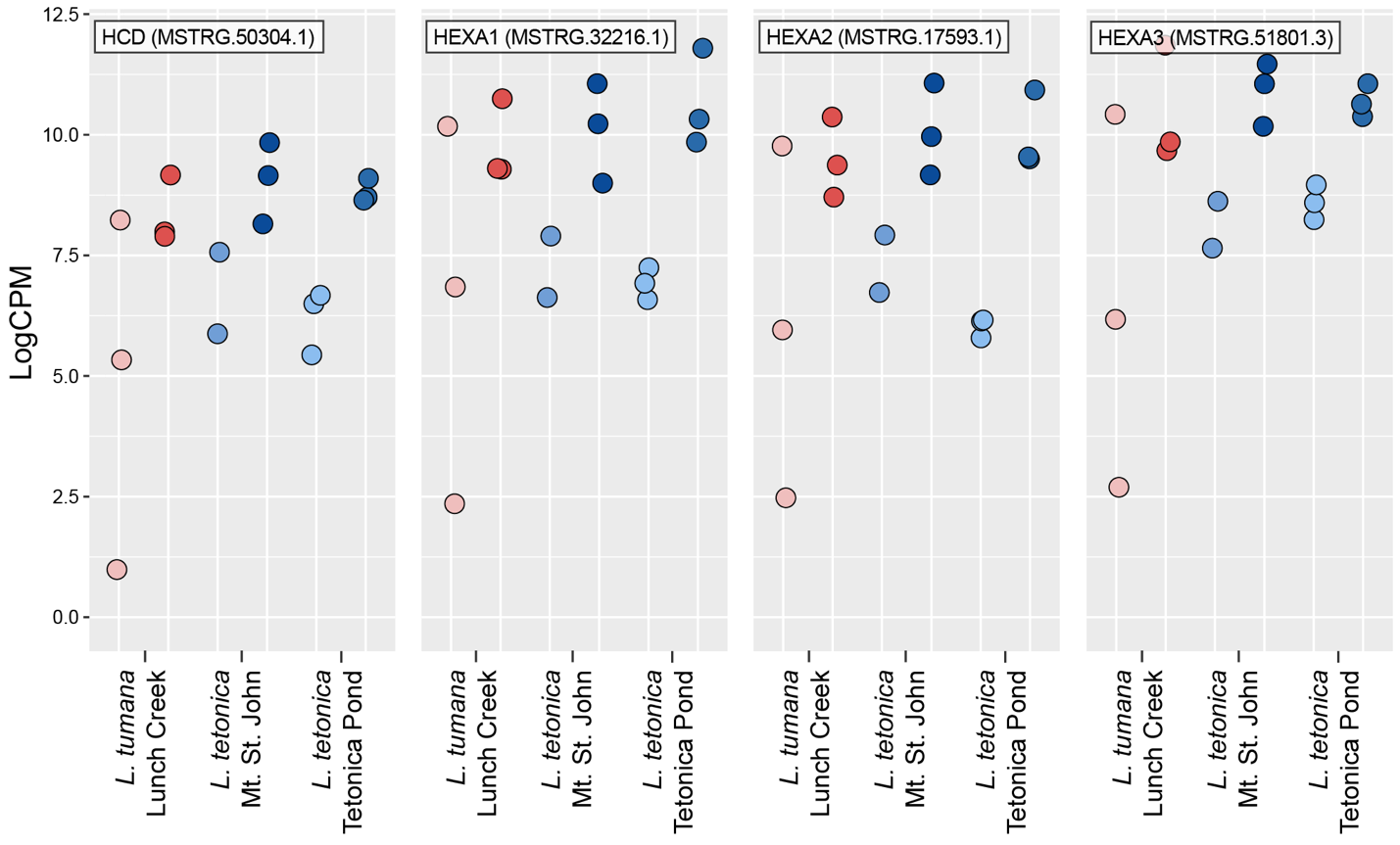

**Figure S9.** Gene expression (logCPM) of hemocyanin (*HCD*) and three hexamerin genes (*HEXA*-1, *HEXA*-2, *HEXA*-3) that were differentially expressed in *L. tetonica*. Expression values are shown for each nymph (circles) with control (lighter) and treatment (darker) individuals highlighted for each species/population.

**Supplementary Videos:**

We have uploaded these files separately with the Supplementary Materials.

**Video S1.** Normal *Lednia tetonica* behavior at temperatures well below their CT_MAX_.

**Video S2.** Loss of righting response at the CT_MAX_ of a *Lednia tetonica* nymph. After 10 seconds, the nymph was pulled from the assay and placed in a recovery chamber.
